# Niche and spatial partitioning restrain ecological equivalence among microbes along aquatic redox gradient

**DOI:** 10.1101/2024.08.09.607300

**Authors:** Katarina Kajan, Rasmus Kirkegaard, Petra Pjevac, Sandi Orlić, Maliheh Mehrshad

**Author notes:** These authors contributed equally (Sandi Orlić, Maliheh Mehrshad). Correspondence: Sandi Orlić, Maliheh Mehrshad.

## Abstract

Microbial metabolic capabilities and interactions shape their niche hypervolume that in turn governs their ecological strategies and ecosystem services. In the context of functional redundancy or ecological equivalence, the focus has been on functional guilds in order to bypass the complex challenge faced by niche theory for disentangling the niche hypervolume. However, in some cases this simplification has been at the expense of ignoring the role of individual genotype of each microbe within a functional guild and fails to explain how the diversity within each functional guild is maintained. In this study, we inspect the metabolic profile of metagenome-assembled genomes along the pronounced redox gradient of the water column in an anchialine cave. Bridging neutral theory of biodiversity and biogeography and niche theory, our analysis uses focal metabolic capabilities while also incorporating individuality by looking into background metabolic capabilities of each individual and further includes spatial distribution of microbes to delineate their niche space. Our results emphasize that differences in background metabolic capabilities are critical for furnishing the niche hypervolume of microbes carrying the same focal metabolic capability and refute their ecological equivalence with their spatial distribution further enables niche partitioning among them.

## Introduction

Defining ecological strategies that individual populations employ in order to ensure their own persistence within the ecosystem (i.e., niche space) is critical for understanding how microbial communities assemble^1–4^. However, there are still fundamental debates regarding the theoretical frameworks and approaches used to address this important conundrum^5–10^. The combination of all biotic and abiotic traits and features that are required for the persistence of each population in the ecosystem build the multi-dimensional hypervolume that is their Hutchinsonian niche^11–13^. While niche theory considers the genomic potential of each population and their niche hypervolume^14^, the highly debated neutral theory of biodiversity and biogeography^15^ collapses groups carrying out focal metabolic processes in functional guilds and considers them as equivalents, thus largely disregarding their individuality^16^. Nevertheless, in both frameworks, the realized metabolic capabilities of organisms are the point of departure for defining their niche space^14^. In recent years several studies have focused on pointing out pitfalls^7^ and settling the debate between existing frameworks^17,18^ or fine tuning them^19^. Recent attempts to utilize high throughput metagenomic approaches to address this topic have further fueled debates on the complexity and validity of determining and disentangling potential versus realized niches from genomic data^5,14^. Consequently, despite being an instrumental concept, niche space remains enigmatic and neither of the existing conceptual frameworks provides a practical approach to resolve this conundrum.

Metabolic capabilities of bacteria and archaea predicted from their genomic content are an important proxy for mapping their niche space at a community scale^1^. Yet, it is important to keep in mind that not all metabolic capabilities encoded in genomes contribute to the realized phenotype of microbes^20–22^. Thus, metabolic similarities or differences between populations should be important for their fitness in order to be considered as a part of their niche hypervolume^14^. However, metabolic capabilities encoded by each population can bring desired ecological strategies into play once they are needed for survival^23,24^. This is particularly relevant for dynamic ecosystems that experience temporal variations in abiotic/biotic conditions^25–29^. Thus, disentangling different dimensions of niches occupied by different populations (via their encoded metabolic capabilities) carries the potential to illuminate the contribution of these populations to ecosystem functioning.

Ecosystem-wide metagenomic studies at genome resolution have become increasingly feasible in the last couple of decades allowing us to comparatively analyze metabolic profiles of community members from different ecosystems^30,31^. Anchialine ecosystems are tidally-influenced subterranean estuaries^32^ that receive input from both seawater and precipitation (i.e., freshwater)^33,34^. The Sarcophagus anchialine cave, studied here, displays multiple physiochemical gradients (strong oxygen and salinity gradients, as well as slight gradients of temperature and pH) causing the strong stratification of the water column and its’ inhabiting microbial community. The pronounced redox gradient developed along the water column in the Sarcophagus cave thus clearly imposes a selection pressure on the community members. Here, we use genome-resolved metagenomics to comparatively study the metabolic profile of community members across the redox potential gradient in different layers of the caves’ stratified water column. In our analyses, we employ aspects of both neutral and niche theory frameworks and utilize a novel practical approach leveraging focal metabolic capabilities (inspired by neutral theory of biodiversity and biogeography) while incorporating background metabolic capabilities of individual populations (following niche theory) as well as their spatiotemporal dynamics. We hypothesize that the genomically encoded metabolic capabilities for energy conservation in different populations contribute to their fitness and thus can potentially become a focal component of their metabolism and a critical dimension of their niche. However, observed differences in the individual metabolic profile of MAGs sharing this niche dimension (i.e., carrying genes encoding the focal metabolic capabilities) preclude their ecological equivalence. Our work contrasts and uses concepts from both the neutral and niche theory to define the niche space of different microbes in the ecosystem and further emphasizes the importance of individuality. In other words, we show that the individual metabolic capabilities of each population are critical for realizing a niche around shared dimensions (i.e. the shared focal function within a functional guild).

## Results and discussion

### Microbial community of the Sarcophagus anchialine cave is strongly stratified

The Sarcophagus anchialine cave studied here exhibits a stable oxygen gradient affecting the community composition along the depth that is reminiscent of community sorting due to the redox gradient within a classical Winogradsky column^35,36^. Furthermore, the salinity gradient contributes to the emergence of specific niches in this cave. While the top layer is practically freshwater, along the depth salinity increases and finally reaches the salinity of marine ecosystems in deeper layers (**Supplementary Fig.1** shows the cave location and vertical profile of different parameters). Physical parameters show similar trends at all three sampling time points (i.e., March, April and August 2021). The halocline and oxycline developed at 2 to 3 meters water depth depending on the month, and hypoxia occurred from 3 m depth of the water column until the bottom of the cave at 12-13 m depth (including sampling points SC3, SC4, SC5, and SC6; **Supplementary Fig.1**).

The pH levels declined sharply within the first 50 cm of depth, consistently remaining within the range of 7.0 to 7.3 below that and throughout hypoxic depths (**Supplementary Fig.1**). Temperature fluctuations exhibited minimal variation, showing a gradual increase with depth in March and April, while in August, the freshwater layer reached a peak of 15.8°C (**Supplementary Fig.1**).

Additionally, the stratification is detected in the concentration of chemicals along the cave profile in April and August measurements. Measured concentrations of HS^-^ peaked below the halocline, reaching a maximum of 13,393 µg/L in April (SC4). Sulfate concentrations increased with depth (April and August) and reached the highest levels in August at the depth of 12 m, measuring 3,200 mg/L (chemical measurements are shown in **Supplementary Table 1**).

A preliminary overview of the microbial community by 16S rRNA gene amplicon analysis shows the temporal development and stabilization of the oxycline from March to August (**Fig.1A**). A higher percentage of ASVs were classified as bacterial in the upper, oxic water layers (>80%; SC1, SC2), whereas archaea dominated in the hypoxic deeper layers (60-78%; SC3 to SC6). The majority of the 5,230 detected ASVs were present only in deeper layers, with 60% assigned to 11 different archaeal phyla (**Supplementary Table 2**). In general, a higher alpha-diversity is recorded in the deeper samples (SC3, SC4, SC5, SC6) as compared to the surface layers (**Fig.1B&C**). Prior to the stabilization of the oxy- and thermocline, Bacteroidota (*Pseudarcicella*) dominated the freshwater layers with a bloom of a single ASV in April (19%) (**Supplementary Fig.2**; **Supplementary Table 3**). By stabilization of the oxycline and thermocline in August, the community of this layer experiences a bloom of Campylobacterota, comprising 7 ASVs, with a single Arcobacteraceae ASV accounting for *ca.* 42% of the community, along with Sulfurimonadaceae (*Sulfurimonas*, *Sulfurovum*, and *Sulfuricurvum*) cumulatively accounting for 59% of the community. Above the stable clines, individual ASV of *Nitrosarchaeum* dominated, making up 13% of the community. In the hypoxic layers, Campylobacterales continued to dominate with Arcobacteraceae at 4 to 7 m depth, but *Sulfurovum* and *Sulfurimonas* in the layers below 7 m (**Supplementary Table 3**). This strong stratification of the microbial community along the depth profile of the cave underlines the importance of physicochemical gradients for structuring the community^37–39^.

**Fig. 1.**
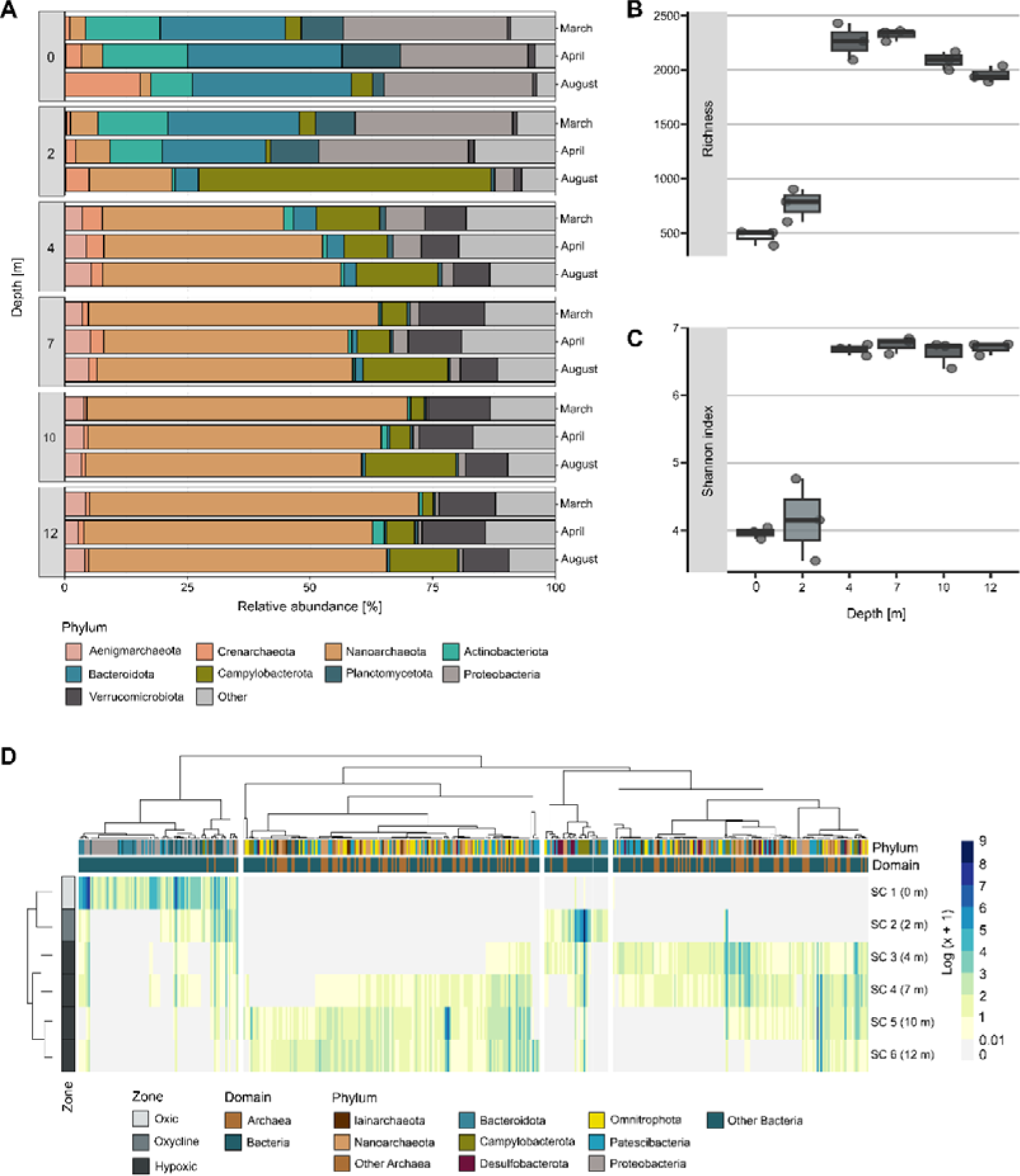
Dynamics and taxonomic diversity of archaeal and bacterial phyla along pronounced environmental gradients in an anchialine cave. **A** Relative abundance assessed by 16S rRNA gene amplicon analysis in three sampling time points (March, April and August). Phyla accounting for less than 5% in relative abundance are categorized as ‘Other’. **B, C** The mean alpha diversity encompassing richness and Shannon index of microbial community along the depth. Boxplots represent the 1st and 3rd quartiles, the line represents the median, and the points are the established data. **D** Vertical distribution trajectory of metagenomic abundances across depth. The cumulative abundances of MAGs were log-transformed (log (x+1)). The top rows are color-coded based on the phylum and Domain of the MAGs. The left column is color-coded according to the vertical distribution of dissolved oxygen. The dendrogram at the top shows the primary clustering of prokaryotic communities based on their prevalence across sampling depths.

To gain a genome-resolved view of the metabolic profile of this stratified microbial community we generated metagenomic datasets from the depth profile samples collected in August, when the oxycline was stabilized. Applying automated binning and dereplicating bins at 95% average nucleotide identity resulted in a total of 418 MAGs (CheckM completeness >=50%, contamination <=10%), of which 119 were high-quality drafts (CheckM completeness >=90%, contamination <=5%). MAGs were assigned to 35 phyla, where most reconstructed MAGs were affiliated to Patescibacteria (79 MAGs; 18.9%), Nanoarchaeota (74 MAGs; 17.7%), Omnitrophota (59 MAGs; 14.1%), and Proteobacteria (49 MAGs; 11.7%) (statistics of reconstructed MAGs and their taxonomy in **Supplementary Table 4**).

Datasets originating from different depths along the cave’s water column represent a strong stratification in their community composition, corroborating the 16S rRNA gene amplicon results (**Supplementary Fig.3**). The uppermost layer hosts a hallmark epilimnetic freshwater community, dominated by *Limnohabitans* and Nanopelagicales^40,41^ (**Fig.1D**). Around 8% of reconstructed MAGs (accounting for *ca.* 35% of MAGs present in SC1) are exclusively present in this stratum and were not detected in other sampled depths (**Supplementary Fig.4**).

The second sampled layer (SC2), coinciding with the thermocline and oxycline, represents a slightly higher alpha diversity compared to the top layer based on ASV analysis (**Fig.1B&C**). Relative MAG abundances confirmed the 16S rRNA gene ASV abundance analysis showing a bloom of sulfur-oxidizing bacteria affiliated to families Arcobacteraceae, Sulfurimonadaceae, and Thiobacillaceae in this layer (**Fig.1D**).

SC2 contained 9 MAGs (2% of reconstructed MAGs, accounting for ca. 11% of MAGs present in SC2) exclusively present in this layer, belonging to taxa Bacteroidales (F082, Paludibacteraceae, UBA12481 and UBA1556), Campylobacterota (Sulfurimonadaceae), Chloroflexota (EnvOPS12), Patescibacteria (BM507) and Proteobacteria (Magnetaquicoccaceae).

The hypoxic lower layers (SC3 to SC6) showed a higher overlap in their community composition apart from 2, 1, and 3 MAGs that were exclusively present in SC3 (Omnitrophota (Koll11) and Patescibacteria (Paceibacterales)), SC5 (Proteobacteria (Alteromonas)), and SC6 (Patescibacteria (two representatives of family UBA2163 and GCA-2747515)), respectively. Along the depth in these 4 datasets, the community gradually shifted with 57 MAGs shared between SC3 and SC4, 92 MAGs shared between SC4, SC5, and SC6, 37 MAGs shared between SC5 and SC6, and 63 MAGs that are shared across all these 4 datasets (**Fig.1D**, **Supplementary Fig.4**). The majority of MAGs present in these four layers (83%) were affiliated to Archaea (38%), Patescibacteria (24%) and Omnitrophota (21%) (**Fig.1D**).

There was only a small overlap in the community of all depths across the redox potential gradient represented by a total of 11 representative MAGs (3% of all reconstructed MAGs) that were affiliated to Nitrosoarchaeum (Thermoproteota), Nanopelagicales (S36-B12), Ilumatobacteraceae (BACL27), Bacteroidales (UBA6244 sp002440885), Arcobacteraceae (CAIJNA01), Phycisphaerales (F1-60-MAGs104), Burkholderiales (Gallionella, Limnohabitans, Limnohabitans_A, and Thiobacillaceae) and Diplorickettsiales (Rickettsiella_A).

### Combined metabolic profile of the community does not mirror the strong stratification of the community composition

While the community composition varied considerably among different depths (sharing only 3% of MAGs, n=11, **Supplementary Fig.5**), the overall community level metabolic profile in each depth seemed to largely overlap (**Supplementary Fig.5**). From 7535 KEGG KO annotated genes, 4911 (ca. 65%) were detected in all 6 depths, without taking into account the taxonomy of the MAGs encoding them (**Supplementary Fig.5**).

Only 186 KEGG KO were exclusively present in the metabolic profile of the SC1 community (**Supplementary Table 5**). While 8% of MAGs were exclusively present in this depth, they contributed only a minor metabolic novelty (2% of all annotated KOs) to the community. These 186 KOs encoded for functions related to KEGG categories metabolism, genetic information processing, and signaling and cellular processes (list of KEGG KOs in **Supplementary Table 6**).

Comparing the profile of annotated genes in each reconstructed MAGs showed a clear clustering that differentiates between Bacteria and Archaea (**Supplementary Fig.6A**). Based on PCoA analyses; at higher taxonomic resolution; the metabolic profile of Thermoproteota representatives seemed to be slightly divergent from other Archaea (**Supplementary Fig.6B**), and a different metabolic profile is evident for representatives of Patescibacteria and Omnitrophota as compared to other bacterial taxa (**Supplementary Fig.6C**).

The observed contradiction where highly stratified communities (only 3% of MAGs shared among all 6 samples) along the cave’s depth profile encode for a highly similar metabolic profiles at community level (65% shared across all 6 samples) is paradoxical. This raises the question whether unique metabolic capabilities of individual populations (i. e., representative MAGs) affect their fitness in occupying different niches and play a role in community assembly or are rather a byproduct of community assembly where functionally redundant microbes can coexist in the available niches.

To address this paradox, we initially differentiated between the common genes (i.e., present in higher abundance among the community) and the rare genes that are most likely involved in environmental adaptations and metabolic specialisation^42–44^. As a proxy, cumulative abundance of MAGs containing each annotated KO was assigned as the abundance of that KO at each depth (**Supplementary data 1**). We then separated the annotated KOs to common and rare based on the sum of their abundance in all depths (threshold of 2000 cumulative abundance; **Supplementary data 2**). Taking the abundance of each KO into account using this approach reveals differences in the prevalence of functions at each depth (**Supplementary Fig.7A&B**). Ordination of the abundance-corrected metabolic profile of each depth based on their dissimilarity for all KOs is mainly driven by differential abundance of MAGs at each depth (**Supplementary Fig.7A&B** and **8A&B**). However, separating the common versus rare KOs in each depth changes the view and highlights a slightly higher dissimilarity between the communities based on rare KOs (**Supplementary Fig.8B&C**). Interestingly, this is more evident in the deeper strata between SC3 and SC4 (13.6% shared MAGs, representing 26.3% and 18.6 % of the communities, respectively) as well as SC5 and SC6 (8.8% shared MAGs, representing 14.5% and 17.1% of the communities, respectively) where the communities are more overlapping. However, this could potentially be affected by the more pronounced dissimilarity of abundance-corrected profiles of these deeper strata from the surface samples (SC1 and SC2) rather than their similarity to each other per se (**Supplementary data 3**). The common KOs seem to have a lower contribution to the dissimilarity of the community’s metabolic profile even after correcting for the abundance of MAGs containing them at each depth (**Supplementary Fig.8B**).

Finally, analyzing the metabolic profile at community level seems inefficient in resolving the paradox of the highly similar metabolic profile encoded by strikingly stratified communities along the depth. Consequently comparing the community metabolic profile does not resolve the impact of niche space on the community composition. Correcting the metabolic profile for the abundance of each KO in the community, and further separating these KOs into common and rare categories gives a marginally better insight into the potential role of rare genes in sustaining dissimilarity among communities. Nevertheless, the signal in the abundance-corrected metabolic profile remains too diluted to allow for further understanding of the role of metabolic genes in occupying certain niches along environmental gradients.

### Focal metabolic capabilities are shared between metabolically divergent clusters

To resolve the role of metabolic capabilities in occupying niches available along the pronounced redox gradients in this cave, we focused on genes involved in focal metabolic components of a certain niche. We selected metabolic pathways of sulfur oxidation, sulfate reduction, and nitrogen fixation for this analysis since they are known to correspond to redox gradients^45–47^.

Inspecting the distribution of KOs along the depth profile (**Supplementary Fig.7A&B**; **Supplementary Table 6**) showed that these metabolic pathways are more prevalent at certain depths (all KOs in these pathways are among the common KOs except for those corresponding to aprA and aprB, which are among rare ones).

Abundance-corrected profiles of KOs showed the peak abundance of functions related to thiosulfate oxidation by the SOX complex and nitrogen fixation at the SC2 layer (**Supplementary Fig.7A**). All genes involved in these functional modules were encoded in the MAG JMF-2110-14-0014.bin.62 (affiliated to Campylobacterota) which bloomed at the SC2 stratum that coincides with the oxycline (**Fig.1D**). Based on MAG abundances, JMF-2110-14-0014.bin.62 was detected as present in all depths and represents the most abundant MAG in layers SC2, SC3 and SC4. Distribution profile of the ASV_p2t_128 (affiliated to Campylobacterota, Campylobacteria_ Arcobacteracea) corroborated this finding. However, it remains unclear if this bacterium is active in the hypoxic layers (SC3 and SC4), or its observed abundance is due to sedimentation and dispersion of the intense bloom developed at SC2. In the Sarcophagus cave microbiome, genes encoding thiosulfate oxidation by the SOX pathway were annotated in 20 MAGs (containing key genes and at least 80% of genes necessary for the complete multienzyme complex; **Supplementary Table 7&8&9**). The MAGs belonged to phyla Campylobacterota (n=7) and Proteobacteria (n=13). PCoA ordination of their annotated metabolic profile separated these MAGs in 4 different clusters (0.6 height as threshold, Elbow method; PERMANOVA; R2 = 0.53, p value = 0.001; **Fig.2A&D**; **Supplementary data 4**) which partially correlates with their taxonomic affiliation (**Supplementary Fig.9&10**; **Supplementary data 5** for statistical significance of the clustering).

**Fig. 2.**
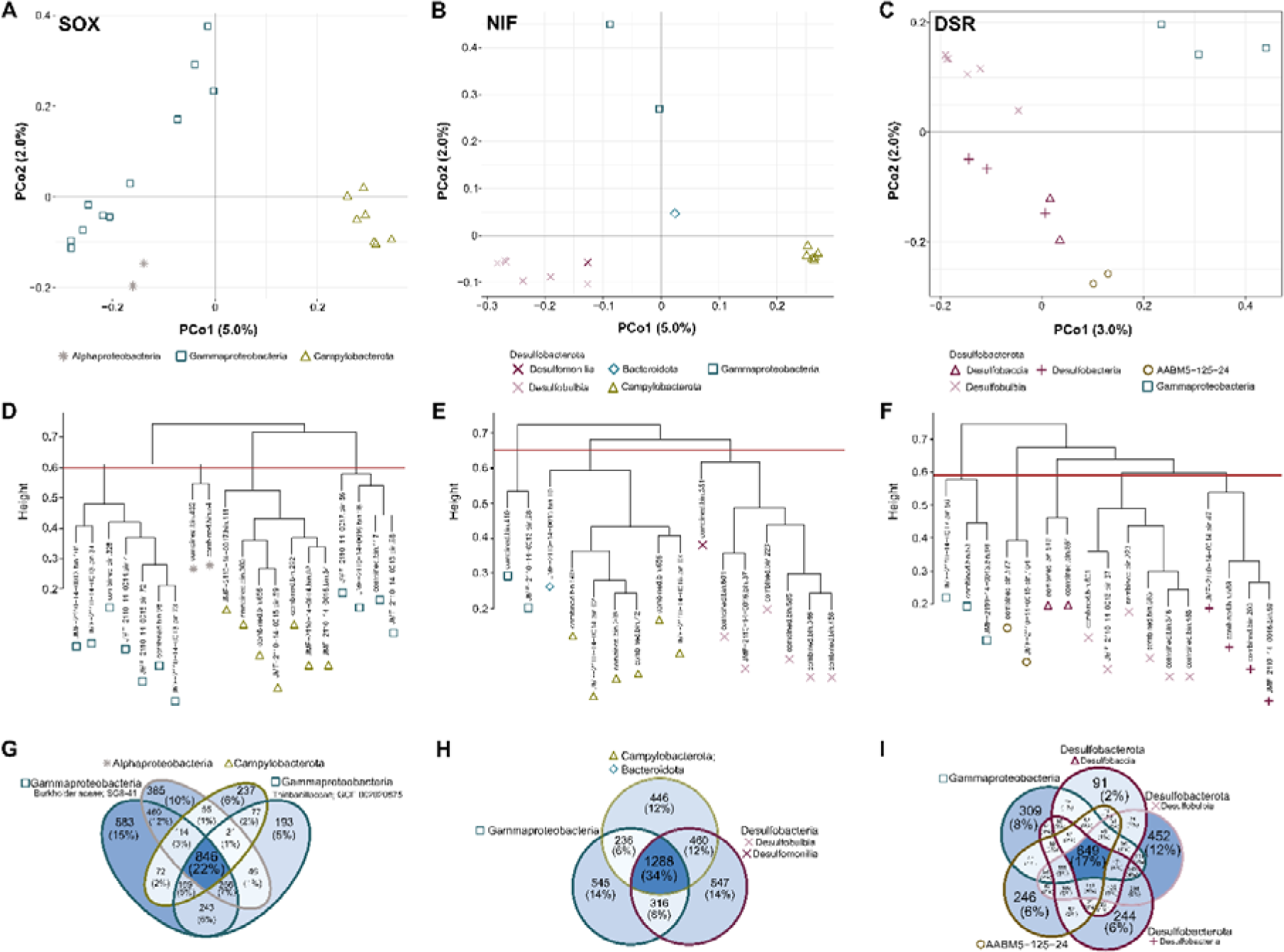
In-depth functional analysis of the microbial metabolic capabilities focusing on sulfur oxidation (SOX), nitrogen fixation (NIF), and sulfate reduction (DSR). Microbial genomes were manually screened for genes associated with prominent metabolic modules. **A, B, C** Diversity of functional capabilities of MAGs encoding each pathway. **D, E, F** Hierarchical relationship between the MAGs and clustering patterns based on their functional capabilities. The red line indicates the operational number of clusters. **G, H, I** Distribution of KOs across diverse MAG clusters. Venn diagrams show the numbers and percentages of unique and shared KOs among MAG clusters.

These 4 clusters shared 846 KEGG KOs in their profile showing an overlap of only 22% among all their annotated KOs. Each cluster also showed a different level of functional overlap with other clusters, while each cluster encoded 5 to 15% of unique annotated KOs (**Fig.2G**; **Supplementary Table 10**).

Moreover, while JMF-2110-14-0014.bin.62, combined.bin.656, and JMF-2110-14-0013.bin.98 MAGs are capable of thiosulfate oxidation by SOX they also encoded genes for nitrogen fixation, another focal metabolic capability (genes nifD, nifK, and nifH). In our database of reconstructed MAGs there are 13 other MAGs that also encoded genes for this metabolic module. MAGs containing nitrogen fixation genes belonged to taxa Bacteroidota (n=1), Campylobacterota (n=6), Desulfobacterota (n=7; Desulfobulbia and Desulfomonilia), and Gammaproteobacteria (n=2; Burkhorderiales) (**Supplementary Fig.11&12**; **Supplementary Table 7&8&11**). Clustering them based on similarity of their encoded metabolic profile resulted in 3 clusters (0.65 height at threshold, Elbow method; PERMANOVA; R2 = 0.43, p value = 0.001; **Fig.2B&E**; **Supplementary data 4**) which again partially correlated with their taxonomic affiliation (**Supplementary data 5** for statistical details). These clusters share 1288 KEGG KOs accounting for a 34% overlap in their annotated KOs (**Fig.2H**; **Supplementary Table 12**). The overall metabolic profile of these clusters contained between 12 to 14% of KOs unique to each cluster.

To further add to the complexity of metabolic overlaps within the cave community, 6 MAGs encoded nitrogen fixation and DSR pathways (**Fig.3**). The dissimilatory sulfate reduction (DSR) pathway (defined here by the presence of marker genes *aprAB*, *dsrAB*, and *sat*) was annotated in 17 MAGs assigned to Desulfobacterota (n= 12), AABM5-125-24 (n=2), and Gammaproteobacteria (n=3) (**Supplementary Fig.13&14**; **Supplementary Table 7&8&13**). Comparing the metabolic profile of these MAGs puts them in 5 different clusters (0.59 height at threshold, Elbow method; PERMANOVA; R2 = 0.58, p value = 0.001; **Fig.2C&F**; **Supplementary data 4**) that shared 649 KOs (22 % of all annotated KOs; **Fig.2I**; **Supplementary Table 14**). Whereas, 2 to 12% of KOs were unique to each cluster (**Supplementary Table 15**). Among these MAGs, the *dsrA* gene encoded by three MAGs affiliated to Gammaproteobacteria were the oxidative type based on phylogeny of *dsrA* (**Supplementary Fig.15**) and they all cluster together (**Fig.2F**). Two of these Gammaproteobacteria MAGs encoded the SOX pathways as well (**Fig.3**). Furthermore, the MAG combined.bin.346 (class Desulfobulbia) encoded both the reductive and oxidative DSR pathway.

**Fig. 3.**
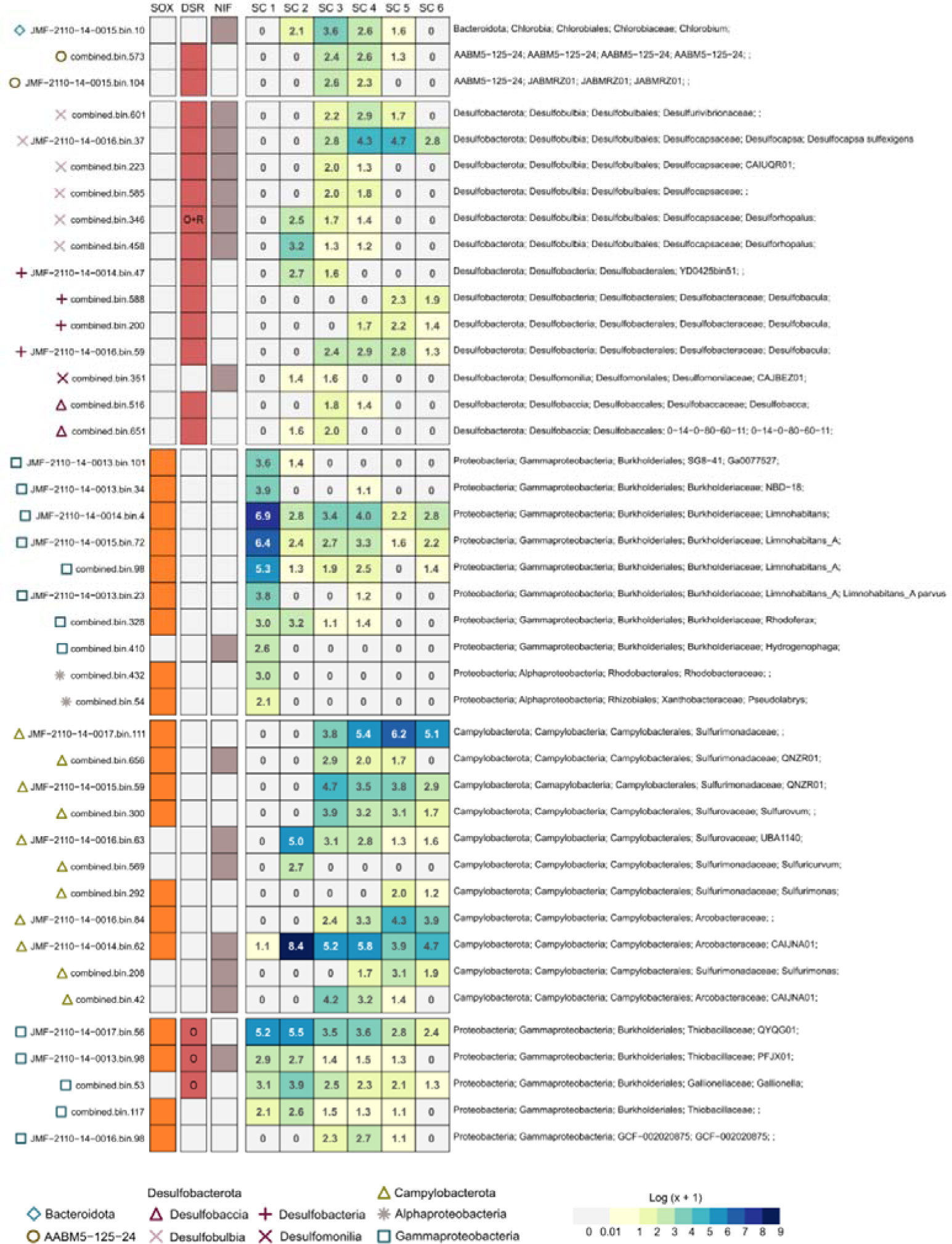
Distribution patterns of MAGs encoding sulfur oxidation (SOX), sulfate reduction (DSR), and nitrogen fixation (NIF) pathways along the vertical gradient. Metabolic analyses of annotated functions separated MAGs containing these functions to five main clusters. The clustering relies on the binary Jaccard distance, quantified using the presence-absence profile of KOs in MAGs. The first three columns indicate the presence of the prominent metabolic module within each MAG. The cumulative abundances of MAGs were log-transformed (log (x+1)).

The metabolic plasticity of MAGs encoding for multiple focal metabolic capabilities and relevance of these functions to redox gradient; leads to an emergent property where all members carrying any of these focal metabolic capabilities (n=42 MAGs) are interconnected within and across communities along the depth profile. They are either directly connected because of carrying multiple focal metabolic capabilities (n=8 MAGs) or because of sharing a niche dimension with other microbes carrying multiple focal metabolic capabilities. This brings in additional biotic dimensions to their niche space either intensifying the competition over resources or leading to niche partitioning via the activity of their background metabolic capabilities or spatiotemporal dynamics^48^. Moreover, metabolic capabilities specifically encoded by each cluster further contribute to their niche hypervolume and rule out their equivalence. If we go beyond the clusters and look at each MAG individually, the level of metabolic overlaps becomes more limited (only 18 KOs are shared among these 42 MAGs; **Supplementary Table 16**). This can be due to high diversity of their background metabolism and to a degree also influenced by MAG incompleteness. Our results highlight how clustering MAGs based on their overall metabolic capability around a specific focal metabolic capability is more informative for deciphering the role of their metabolic capabilities in their niche hypervolume, rather than clumping all MAGs carrying a specific metabolic function together as a guild with assumed functional redundancy.

### Metabolic and spatial niche partitioning enable co-existence of clusters with redundant focal metabolic capabilities

Our results showed that MAGs containing genes for focal metabolic capabilities encoded differing metabolic profiles. Comparing the metabolic profile of these 42 MAGs containing at least one of the studied focal metabolic capabilities separated them in 5 main clusters (PERMANOVA; R2 = 0.06, p value = 0.002; **Supplementary Fig.16A&B**; **Supplementary data 4** for statistics). While these clusters partially correlated with taxonomy and depth distribution, they did not separate based on the focal metabolic capability that each MAG encoded (**Supplementary data 4**). These 5 clusters (each containing multiple MAGs) shared 921 KOs (19% of all KOs annotated in all clusters; **Supplementary Fig.17**; **Supplementary Table 17**). At the MAG level, the number of KOs shared between MAGs of each cluster ranged between 230 to 627 (1342 KOs in each MAG on average). Separately clustering MAGs containing each focal metabolic capability showed that generated clusters shared 864 (individual MAGs in each cluster share 349-989 KOs), 649 (individual MAGs in each cluster share 505-1166 KOs), and 1288 (individual MAGs in each cluster share 323-1111 KOs) KOs for SOX, DSR, and NIF clusters, respectively (**Supplementary Table 18**). As expected, there was a significant negative correlation (Pearson’s correlation; R2 = -0.76, p value = 0.004) between the percentage of total shared KOs and the number of MAGs in each cluster (**Supplementary Fig.18**; **Supplementary Table 18**). This suggests that in order to sustain the higher diversity within each focal metabolism, populations within each cluster should maintain more differences in their background metabolic capabilities. While clustering groups containing a certain focal metabolic capability enabled us to identify MAGs with higher metabolic overlap (snippet from neutral theory), these clusters and all MAGs within each cluster contain metabolic differences from each other. These metabolic differences contribute to niche partitioning and allow these microbes to further furnish their niches around these shared dimensions (in line with niche theory). This shows that MAGs carrying the same focal metabolic potential could not be considered as equivalents with full niche overlap. Meaning that calling such MAGs as functionally redundant or ecological equivalents is not justified and they rather share dimensions within their multidimensional niche hypervolume.

It is important to note that the focal metabolic capabilities discussed here would not be a focal component of all realized niches of microbes carrying them across the redox gradient^37,49–51^. For example, JMF-2110-14-0014.bin.62 is present in all sampled layers. This MAG forms an intense bloom that coincides with the oxycline (SC2 layer) which is an optimal redox condition for energy conservation via SOX, thus justifying the assignment of SOX as its focal metabolic capability. This MAG is just at the threshold of presence in the fully oxic surface layer but is present throughout the hypoxic deep layers (being the most abundant MAG in SC3 and SC4). However, its encoded SOX and NIF metabolic capability might have a different relevance for its realized niche in different layers (e.g., by being coupled to different electron acceptors that is affecting their energy yield). Furthermore, sedimentation of the dense bloom developed at SC2 could contribute to the high prevalence of JMF-2110-14-0014.bin.62 in deeper strata making it more challenging to directly connect its abundance to its metabolism.

In addition to background metabolic capabilities of MAGs belonging to the same or different clusters, their spatial distribution can further contribute to their niche partitioning (**Fig.3**). Different clusters have a different pattern of distribution (**Fig.3**; **Supplementary Fig.19**). The PCoA ordination of the abundance profiles of these 42 MAGs is primarily separating them based on their presence or absence in the surface layer (**Supplementary Fig.20**). Mainly Gammaproteobacteria and Alphaproteobacteria affiliated MAGs had highest abundances in the surface layer SC1, also including representatives with highest prevalence in SC2 while keeping a high prevalence in SC1. An exception is MAG JMF-2110-14-0016.bin.98 that is forming a cluster together with 4 other Gammaproteobacteria MAGs based on their metabolic profile (**Supplementary Fig.16**) but has a different spatial distribution pattern and is absent in SC1 and SC2 (**Fig.3**). This MAG carries SOX and the complete pathway for dissimilatory nitrate reduction, which could be coupled in the hypoxic layers. Among the other 4 MAGs within this same metabolic cluster only JMF-2110-14-0013.bin.98 also encodes SOX, dissimilatory nitrate reduction, and oxidative DSR which had its peak abundance in the surface.

MAGs that were absent in the surface layer were represented in 3 clusters (including bloom forming JMF-2110-14-0014.bin.62 that is present in SC1 but only at the threshold of our cutoff for presence, **Fig.3**). Within each cluster however, the range of distribution across different depths varied (**Fig.3**; **Supplementary Fig.9-14**) further alleviating the competition around this niche dimension at a spatial level. Specifically, MAGs encoding DSR belonging to the phyla Desulfobacterota and AABM5-125-24 were separated in 4 clusters based on their metabolic profile and were all absent in the surface layer. Within the cluster affiliated to class Desulfobacteria, 3 MAGs in the genus Desulfobacula (family Desulfobacteraceae) were prevalent in the deeper strata (different range of prevalence across SC3 to SC6) whereas the remaining MAG (JMF-2110-14-0014.bin.47, affiliated to family YD0425bin51) was only present in SC2 and SC3. Interestingly, the ordination of metabolic profiles (**Fig.2C**) placed this MAG close to the 2 MAGs in the Desulfobaccia cluster that had their highest prevalence in SC3 (with expansion to either SC2 or SC4). This could hint that the metabolic capabilities of this MAG are close to other MAGs prevalent in the same strata due to niches defined at this layer with its specific physicochemical features. While spatial niche partitioning was more pronounced for this MAG, the 3 Desulfobacula MAGs had only slight differences in their range of presence and coincide in SC5 and SC6. Nevertheless, while these MAGs (completeness higher than 97%) shared 1166 KOs this accounts only for 68, 71, and 89% (in MAGs combined.bin.200, JMF-2110-14-0016.bin.59, and combined.bin.588, respectively) of their annotated KOs (**Supplementary Table 19**). The remaining 11 to 32% of the unique KOs that each of these MAGs carry could potentially contribute to further furnishing their niche and enabling them to coexist in the same water strata. Our results highlight the importance of background metabolic capabilities (**Fig.4A**, **Supplementary Table 20**) in the niche space of lineages encoding focal metabolisms (**Fig.4B&C**) with overlapping or divergent spatial distribution (**Fig.3**). Encoded KOs outside the focal metabolic capabilities enable different lineages to shape shift and partition their niches around the shared dimensions within and across different lineages further rebutting their ecological equivalence (**Fig.4D**, **Supplementary Table 20**).

**Fig. 4.**
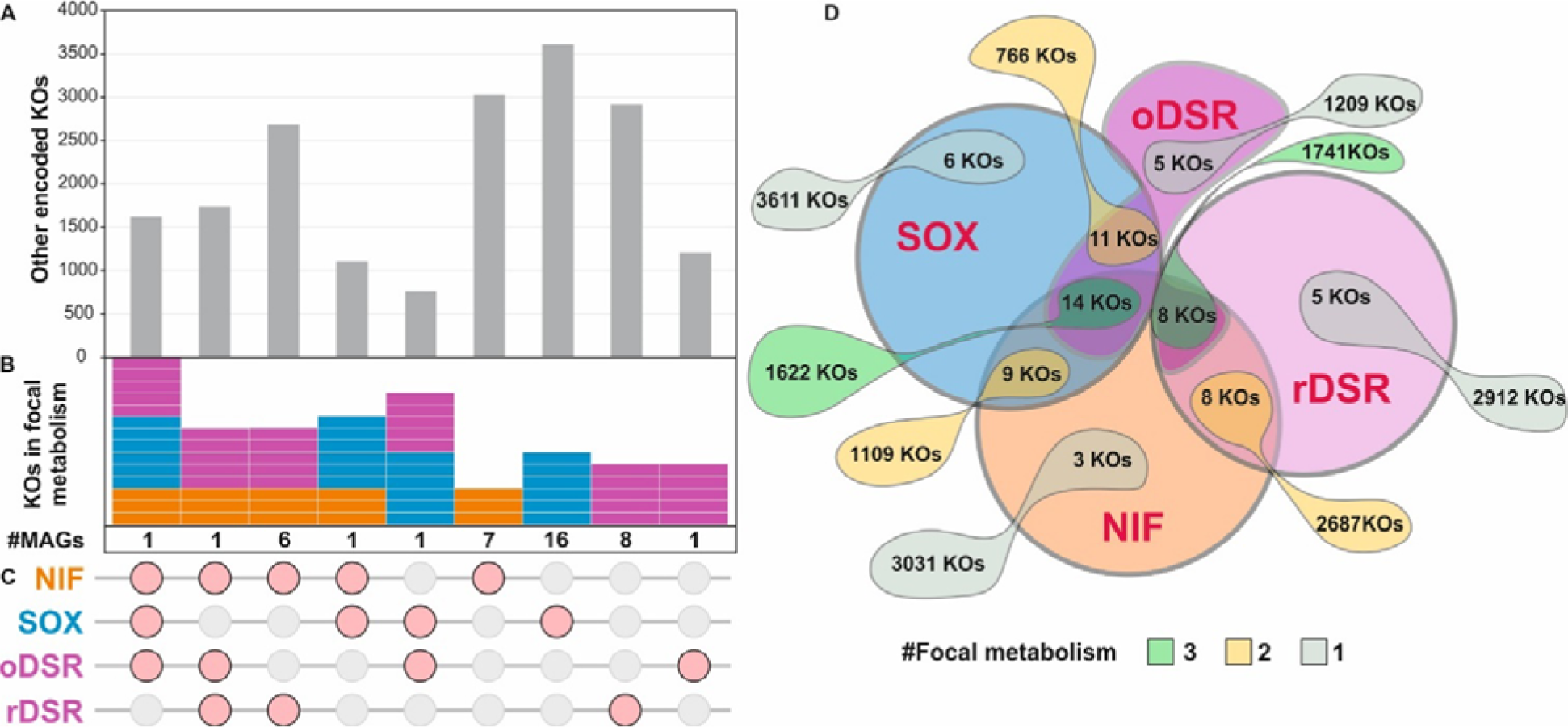
Significance of background metabolic capabilities for niche partitioning. **A, B, C** up set plot showing the overlap of focal metabolic capabilities among MAGs carrying each function and their background metabolic capability represented as all unique KEGG KOs encoded in MAGs of each category. **D** Barba-plot (inspired by characters of Barbapapa animation^1^ that are able to shapeshift at will) shows the extent of background metabolic genes in a simplified depiction of the complexity of niche hypervolumes around the shared niche dimensions for focal metabolic capabilities.

## Conclusion

The capability to carry out the same metabolic function^52^ or carrying it out at the same rate^53^ have been used to assign lineages as functionally redundant in the environment^54^. From the viewpoint of securing ecosystem services and assessing the impact of diversity loss on the provision of certain services, the ability of the community to buffer those disturbances via functional redundancy seems critical. However, species capable of carrying out the same metabolic capability and provisioning the same service after the disturbance must have been adaptable to these disturbances, which is already refuting their equivalence with their less successful counterpart. Additionally, the still debated functional redundancy concept has spilled over to assessing the basics of community assembly and their eco-evolutionary dynamics^53,55,56^. Functional redundancy or equivalence has been discussed at the level of ecosystem, individual or metabolic modules^52,57^. In the highly dynamic community of microbes, specifically those developed along strongly pronounced physicochemical gradients; it seems premature to presume that carrying certain shared metabolic functions will be the basis for equivalence and redundancy, without considering other dimensions of organisms’ niche hypervolume.

While focal metabolic functions are facilitated by a number of microbes, the metabolic profile of each microbe (i.e., a unit of evolutionary selection) is unique. This slightly different profile could give each organism an edge to defy the limitations imposed by functional redundancy and maintain a unique niche around this shared dimension. Thus, the answer to what defines the niche space lies in the differences rather than similarities, as we clearly detect representatives of different species carrying the same focal metabolic capability that are prevalent in the same water layer, thus co-existing yet not redundant.

Encoded metabolic capabilities are not directly linked to the realized phenotype; factors involved in niche partitioning cannot be solely explained by the metabolic profile of microbial lineages and their spatial dynamics. In addition to drift, spatiotemporal dynamics, and transcription control that play a role in occupying niches by different lineages, niche partitioning can be imposed via either symbiotic or parasitic/predation interactions. Different intensity of host-specific top-down controls can play a role in alleviating the competition for the non-target occupant of these niches. All these factors further contribute to niche partitioning and add to the complexity of the niche hypervolume, however, due to existing limitations in resolving them in a system-oriented and high-throughput method, our knowledge in this aspect remains limited to our currently obstructed view which is nevertheless wide enough to rebut ecological equivalence among microbes carrying the same focal metabolic capabilities.

## Material and methods

### Sarcophagus Cave site description and sampling schema

The anchialine cave studied here is located in the karstic Upper Cretaceous (Senon) limestone with rare fragments of dolomite in the eastern coastal area of the Adriatic Sea near the Krka River estuary, Croatia (**Supplementary Fig.1**). The carbonate rocks of the karst area contain channels and fissures porosity that store and transfer a significant amount of groundwater of different salinities^32^. The climate in this area is Mediterranean, with dry and hot summers and mild, rainy winters^58^. The mean annual precipitation at the Šibenik meteorological station was reported at 748 mm in 2021 (Croatian Meteorological and Hydrological Service, www.meteo.hr).

The small vertical entrance shaft of the cave is located at about 580m distance from the estuary shore at an elevation of 5m from the sea level. The overall depth of the cave is about 18m, whereas the vertical shaft channel is about 8m (the dry part measures 6m plus 2m of water, depending on the tides). The known depth of the water body is around 12m. The anchialine cave investigated in this study had no visible signs of direct or indirect anthropogenic influence; therefore, it is presumed to be a pristine habitat. However, a stone sarcophagus brought to the southeast of the entrance indicates that throughout history, the cave has been used to feed livestock, most likely goats and sheep.

Anchialine caves on the Adriatic coast^59–62^ are spatially complex habitats and sampling them requires professional divers and speleological equipment. Technical difficulties in sampling limits the possibility of more frequent sampling or collecting larger volumes of water. From our study site, water samples were collected at six depths along the salinity gradient at 0, 2, 4, 7, 10, and 12m depths (SC1, SC2, SC3, SC4, SC5, and SC6) in March (17.03.), April (14.04.) and August (30.08.) 2021. Samples from the first two depths were collected using a Niskin bottle. A professional cave scuba diver collected samples from depths below 3m. All samples were collected in sterile polycarbonate bottles. For the March and April samplings, 2L of water was collected from each depth, whereas for the August sampling, we collected 5L of water from each depth in a Niskin bottle carried by the diver.

### Physical and chemical parameters

Physical parameters (temperature, salinity, conductivity, dissolved oxygen, pH, TDS and chlorophyll) were measured *in situ* along the depth using a Multisensor CTD probe (EXO2, YSI, USA) carried by the diver. Samples for chemical analysis were collected in acid-cleaned LDPE (Nalgene) bottles and stored at - 20°C until analysis, which was performed within one month.

Concentrations of the cations (Ca^2+^, Mg^2+^, Na^+^), anions (Cl^-^, SO_4_^2-^) and dissolved organic carbon, were determined in filtered water samples in the Hydrochemical Laboratory of the Croatian Geological Survey. The concentrations of cations and anions were measured on Dionex ICS-6000 DC (Thermo Fisher Scientific, Waltham, MA, USA). DOC was analyzed using the HACH QBD1200 analyzer.

Nutrients were analyzed using a UV/Vis spectrophotometer (Specord 200 Plus, Analytik Jena, Germany) in unfiltered samples according to Strickland and Parsons (1972)^63^ for the determination of ammonium (NH_4_^+^), phosphate (PO_4_^3-^), nitrite (NO_2_^-^), and silicate (SiO_4_^4-^). Nitrates (NO_3_^-^) were analyzed according to the vanadium reduction protocol^64^.

Samples for reduced sulfur species (RSS) analysis were collected without exposure to oxygen and analyzed within 24 hours. The total reduced sulfur species (RSStot) were analyzed by electrochemical methods at the hanging mercury electrode (663 VA Stand) connected to the potentiostat (PG STAT 128N, Methrom, Netherlands) as previously described^65,66^. Briefly, RSStot accumulation occurs at -0.2 V (vs. Aa/AgCl) on the Hg electrode surface and forms HgS. The reduction of the formed HgS occurs at -0.68 V (vs. Ag/AgCl) under the given experimental conditions when the potential of the Hg electrode is shifted to more negative values. The current of the detected peak is proportional to the concentration of RSStot. To distinguish between volatile and non-volatile RSS, samples are acidified and purged with nitrogen. After restoring the pH, the samples are analyzed as described and the concentration of the non-volatile fraction (RSSnv) is determined.

### Biomass collection and DNA extraction

To collect the biomass, water samples were filtered on 0.22 µm pore-size polycarbonate filters (Whatman Nuclepore Track-Etch Membrane, diam. 47 mm). The flow through was used for chemical characterization, and filters were stored at -20°C until further processing. Total genomic DNA from filters was extracted using the DNeasy PowerWater Kit (Qiagen, Inc., Valencia, CA, USA) following the manufacturer’s instruction.

### 16S rRNA amplicon analysis

A portion of the extracted DNA was used to amplify the hypervariable V4 region of the prokaryotic 16S rRNA gene using primer pair 515F Parada (5′-GTG YCA GCM GCC GCG GTA A -3′)^51^ and 806R Apprill (5′ GGA CTA CNV GGG TWT CTA AT -3′)^67^, modified with two 16 bp sequences which allow for sample barcoding in a second PCR step^68^. As described in detail in Pjevac et al. (2021)^68^, samples were amplified, barcoded, purified, normalized, and prepared for sequencing on an Illumina MiSeq System in paired-end mode (v3 chemistry, 2x 300 bp; Illumina, San Diego, CA, United States) at the Joint Microbiome Facility of the Medical University of Vienna and the University of Vienna. Sequence data were processed in R using DADA2 following the workflow by Callahan et al. (2016)^69^. Amplicon sequence variants (ASVs) were inferred across all samples in pooled mode. Further details on sequence trimming and settings for quality filtering are described in Pjevac et al. (2021)^68^. Taxonomic assignment was done by mapping 16S V4 ASV sequences against the SILVA SSU Ref NR 99 database (v. 138.1). The sequences assigned to prokaryotic taxa with 16S V4 region were kept for further analysis. Prior to analysis, chloroplastic and mitochondrial sequences were removed and the ASV table was rarefied to the same total number of sequences.

### Metagenome sequencing, assembly, and binning

Extracted DNA from the depth profile collected in August (30.08.2021) was used for metagenome sequencing (PE150) using Illumina NovaSeq 6000 (Eurofins Genomics Europe Sequencing GmbH, Germany).

Metagenomic reads were trimmed using cutadapt (v. 2.10)^70^. Trimmed reads were then assembled using megahit (v. 1.1.2)^71^. Assembled contigs shorter than 1 kbp were removed using seqtk (v. 1.3) (github.com/lh3/seqtk).

To bin the assembled contigs metagenomic reads were mapped to the assemblies using minimap2 (v. 2.17)^72^. The calculated read mappings were then converted using samtools (v. 1.11)^73^ for binning using metabat2 (v. 2.15)^74^. To improve the binning outcome, all generated metagenomes were co-assembled and binned into combined MAGs following the same method.

### MAG analyses

Reconstructed MAGs were then quality checked using QUAST (v. 5.0.2)^75^ and CheckM (v. 1.1.1)^76^. A total of 1420 MAGs were generated and only MAGs with completeness >= 50% and contamination =< 10% were used for further analysis. These MAGs were de-replicated at 95% average nucleotide identity that is the boundary for operational definition of species^77–79^. Taxonomic affiliation of these MAGs was then assigned using GTDBtk (v. 1.5.0)^80^.

Abundance of reconstructed MAGs in each metagenome was calculated as the fraction of the total reads mapping to the contigs in a given MAG. These counts were then normalized for the depth of the sequenced metagenomes. The threshold for considering a MAG present in a metagenome was set to a minimum of 2 per depth.

### Functional annotation

Gene prediction on good quality MAGs was done using Prodigal^81^. MAGs were initially annotated using Prokka (v1.12)^82^. Further functional annotation of the predicted proteins was done using eggnog-mapper (v. 2.2.1) with the eggnog_5.0 database^83,84^. The eggnog-mapper output was used to assign enzyme EC numbers, KEGG orthology (KO), Cazy annotations, pathways and modules to the predicted genes. All annotations of key genes were further inspected and manually checked for their conserved domains using NLM’s Conserved Domain Database (CDD) search^85^. The KEGG database was used to assign pathways, brite and modules of KOs^86^.

For each annotated KO within the MAGs, the cumulative abundance of MAGs carrying this gene in each depth was assigned as the abundance of that gene in the microbial community of each depth. The annotated KOs were further classified into common and rare categories, based on the summation of their cumulative abundances across all depths. The determination of the KOs’ cutoff point involved analyzing the frequency histogram distribution of KOs per depth and in total, resulting with a threshold of a cumulative abundance sum of 2000 to differentiate between common and rare KOs.

### Gene phylogeny reconstruction

To confirm the oxidative versus reductive designation of dsrA genes annotated in our reconstructed MAGs, a phylogenetic tree by including reference sequences was reconstructed. All these dsrA protein sequences were aligned using kaling (v2.04)^87^ and their phylogeny was reconstructed using FastTree (v2.1.11)^88^.

### Data analysis and visualization

All statistical analysis and visualization were done in the R environment (version 4.2.2.)^89^ using packages phyloseq^90^, vegan^91^, dplyr^92^, ggplot2^93^, ggVennDiagram^94^ and Pheatmap^95^.

The alpha diversity (richness and Shannon index) of ASVs was estimated using the vegan package. To assess the variation in microbial community composition through amplicon and metagenome analysis across different depths, a principal coordinate analysis (PCoA) was performed. This analysis utilized Bray-Curtis dissimilarity using Hellinger-transformed data and was carried out using the ade4 package^96^. The significance of the difference in community composition between depths was tested with a permutational multivariate analysis of variance (PERMANOVA; adonis2 function, vegan package in R). Cumulative abundances of MAGs and depths were clustered based on the clustering distance “correlation” (Pearson’s correlation) and clustering method “complete” (pheatmap package). To reduce skewness of the cumulative abundances of MAGs, prior to clustering and visualization a logarithmic transformation (log(x+1)) was applied.

To assess the functional variation of MAGs, a PCoA was conducted based on the binary presence-absence of KOs per MAG. This analysis utilized binary Jaccard distance and was carried out using the ade4 package. The cumulative abundances of prevalent and rare KOs per depth were clustered based on the Euclidean distance and clustering method “complete” (pheatmap package). Furthermore, to evaluate the functional variation per depth, nonmetric multidimensional scaling (NMDS) was carried out using the variation of abundance-corrected metabolic profile per depth. This analysis employed Bray-Curtis dissimilarity with Hellinger-transformed data and was performed using the vegan package.

For further functional analysis of the microbial community, three dissimilatory pathways were analyzed: sulfur oxidation (SOX), sulfate reduction (DSR), and nitrogen fixation (NIF). Selected genes within the modules were: SOX (soxA - K17222, soxX - K17223, soxB - K17224, soxC - K17225, soxY - K17226, soxZ - K17227), DSR (aprA - K00394, aprB - K00395, sat - K00958, dsrA - K11180, dsrB - K11181), and NIF (nifD - K02586, nifH - K02588, nifK - K02591). Only MAGs with completeness >70% and contamination <5% that also contained at least 80% of genes involved in the complete module (SOX, DSR or NIF) were used in this analysis. To statistically prove that the incompleteness of MAGs does not affect this analysis, we used the Mantel test (Spearman’s correlation) to correlate the raw and subsampled dataset (**Supplementary data 6**). Datasets were subsampled to the lowest number of KOs per MAG using the “rrarefy” function in the vegan package. Mantel test based on the Euclidean distance showed high correlation between raw and subsampled distance of dataset (SOX: r=0.897, p=0.001; DSR: r=0.8661, p=0.001; NIF: r=0.9684, p=0.001), while significant lower correlation was detected based on the binary Jaccard distance (SOX: r=0.4991, p=0.001; DSR: r=0.3494, p=0.006; NIF: r=0.7503, p=0.001). To distinguish the diversity of MAGs within the selected modules, a PCoA of all MAGs’ KOs was conducted based on the binary Jaccard distance of presence-absence data. The following data was further clustered based on the clustering method “complete”. To determine the operational number of clusters, the “elbow” method was used to assign the location of the cutoff point using the “fviz_nbclust” function (package factoextra^97^). To statistically support the significance of determined clusters, the difference of groups’ diversity was assessed with PERMANOVA (adonis2 function, vegan package in R).

## Data availability

All metagenomes, amplicon data and MAGs were submitted to GenBank and are accessible under the accession number PRJNA1052463. Accession numbers are presented in **Supplementary Table 21** All the material required to reproduce the results of this study are presented in the text and in the supplementary information. All reconstructed MAGs are publicly available via figshare under the link https://figshare.com/s/01354c02b131bc2c44f6.

## Acknowledgement

This research was partially supported under the project STIM–REI, Contract Number: KK.01.1.1.01.0003, a project funded by the European Union through the European Regional Development Fund – the Operational Programme Competitiveness and Cohesion 2014-2020 (KK.01.1.1.01). Maliheh Mehrshad was supported by a grant from the Swedish research council for sustainable development, FORMAS (grant no. 2021-00546). We gratefully acknowledge diver Branko Jalžić from the Croatian Biospeleological Society and Neven Cukrov from the Ruđer Bošković Institute for the logistic support and sample collection. We acknowledge Marija Marguš from Ruđer Bošković Institute and Tamara Marković from Croatian Geological Survey for performing the chemical analyses, Ema Kostešić and Andrea Čačković from Ruđer Bošković Institute for their assistance and valuable help during the field and lab work, Jasmin Schwarz, Gudrun Kohl and Julia Ramesmayer for technical assistance with sample processing in the JMF laboratories. The computations and data handling were partially enabled by resources in project NAISS 2023/22-1001 provided by the National Academic Infrastructure for Supercomputing in Sweden (NAISS) at UPPMAX, funded by the Swedish Research Council through grant agreement no. 2022-06725.

## Author contribution

KK and SO initiated the study. KK led the sample collection and processed samples together with PP. KK, MM, PP, and RK performed bioinformatic analysis. KK and MM interpreted data, developed the discussions, and drafted the manuscript. All authors read and approved the final version of the manuscript.

## Footnotes

1 https://en.wikipedia.org/wiki/Barbapapa

## Supplementary figures

**Supplementary Fig. 1.**
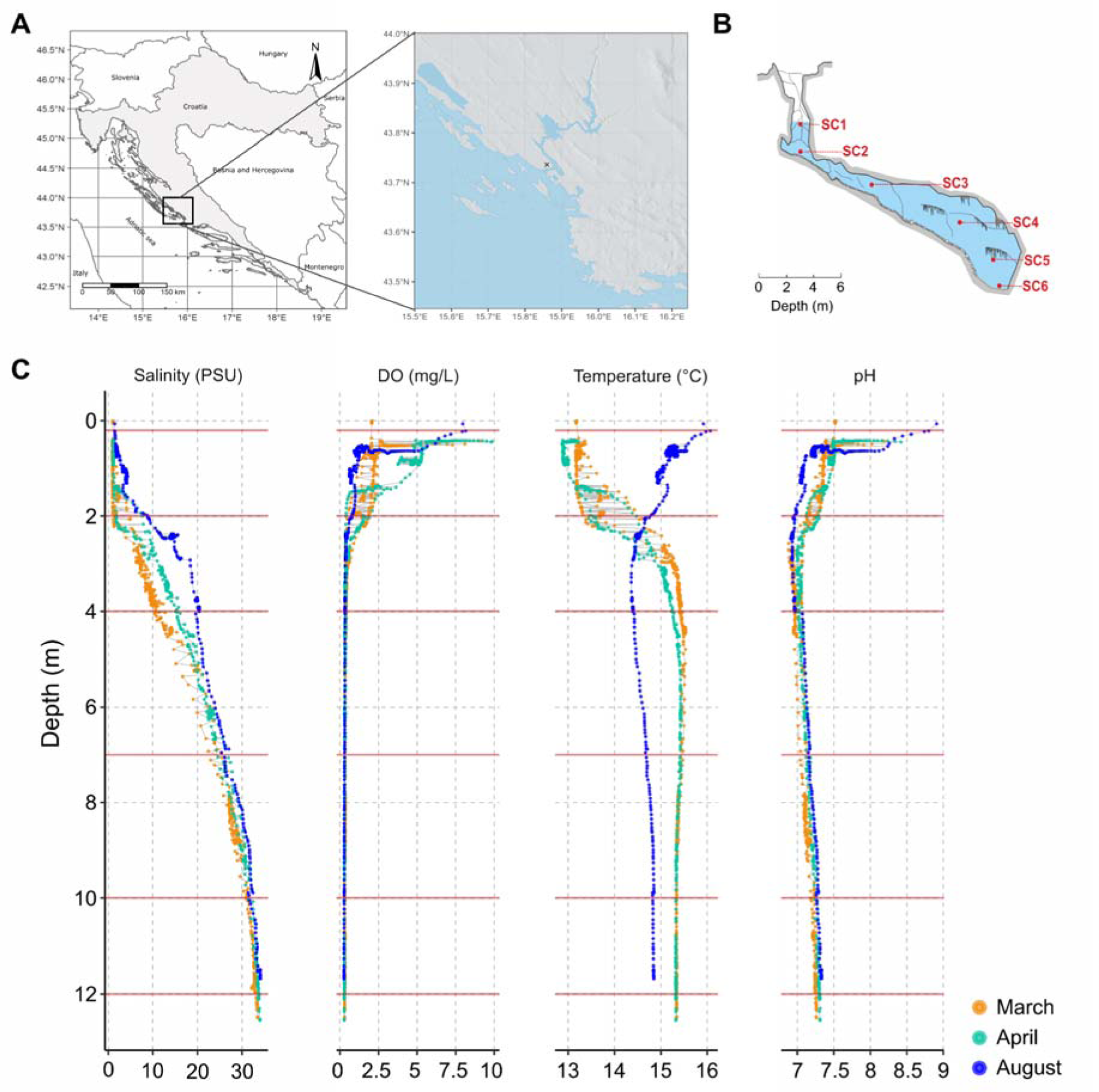
**A, B** Cave location and general vertical structure of the water column in Sarcophagus cave. The study site (indicated by “x”) is located 580 meters inland from the nearest coast. **C** Physicochemical profiles acquired using a multi-parameter data sonde showing salinity, dissolved oxygen (DO), temperature and pH across a depth profile in three sampling time points (March, April and August).

**Supplementary Fig. 2.**
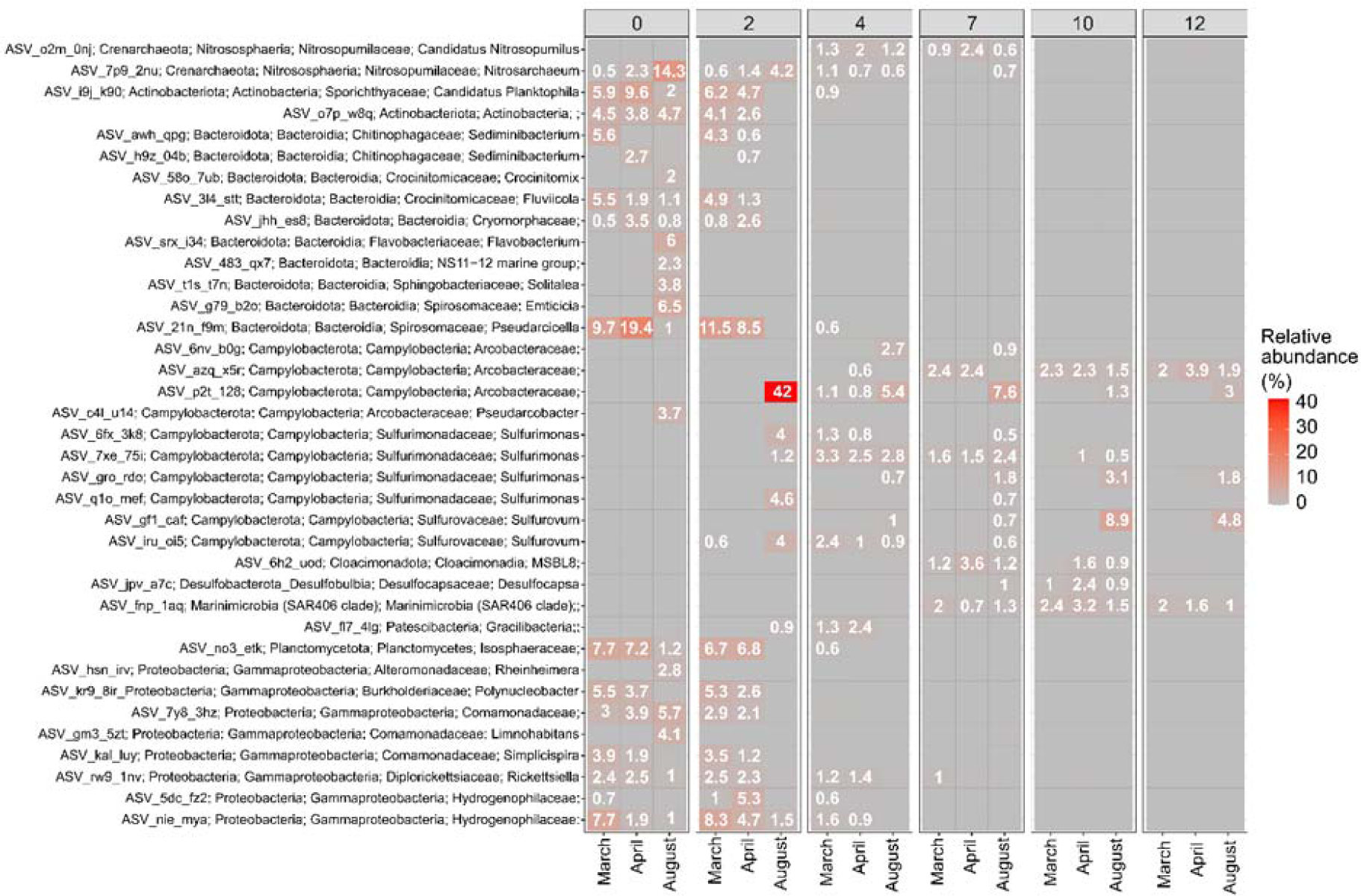
Heatmap depicting the most abundant ASVs in Sarcophagus cave across varying depths and months determined through 16S rRNA gene amplicon analysis.

**Supplementary Fig. 3.**
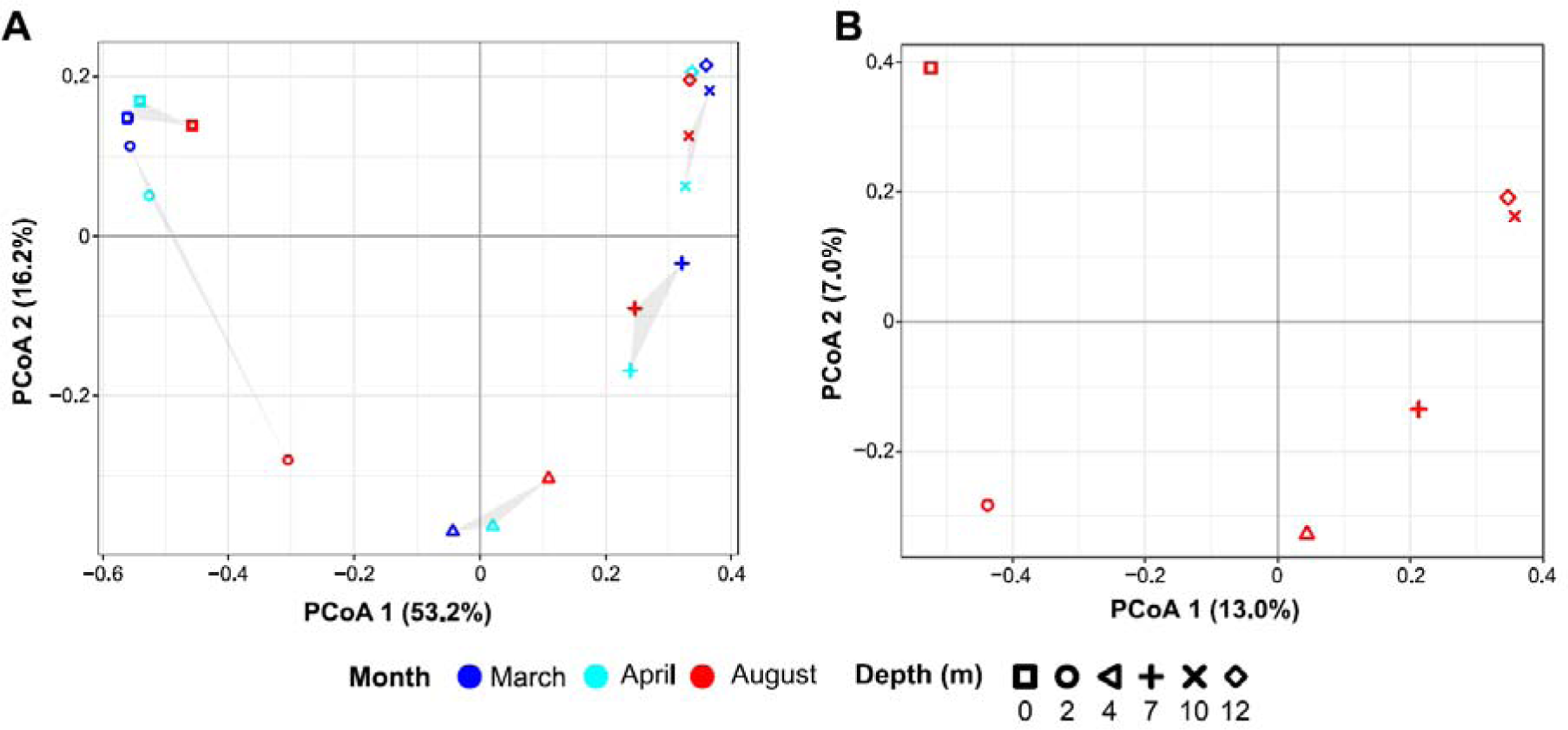
Diversity of Sarcophagus microbial community based on **A** 16S rRNA gene amplicon analysis, and **B** metagenomic analysis.

**Supplementary Fig. 4.**
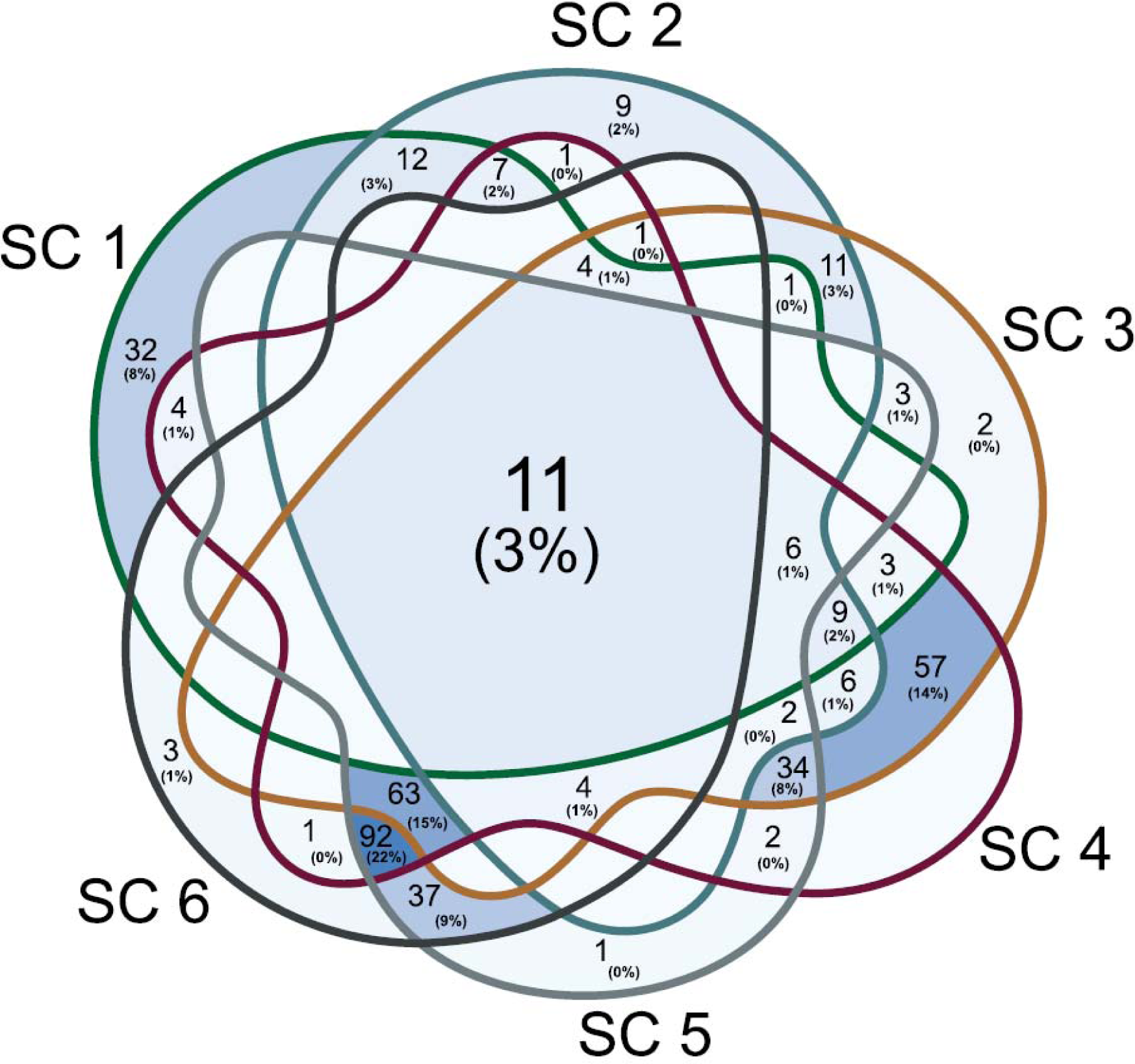
Venn diagram showing unique and shared MAGs at different depths.

**Supplementary Fig. 5.**
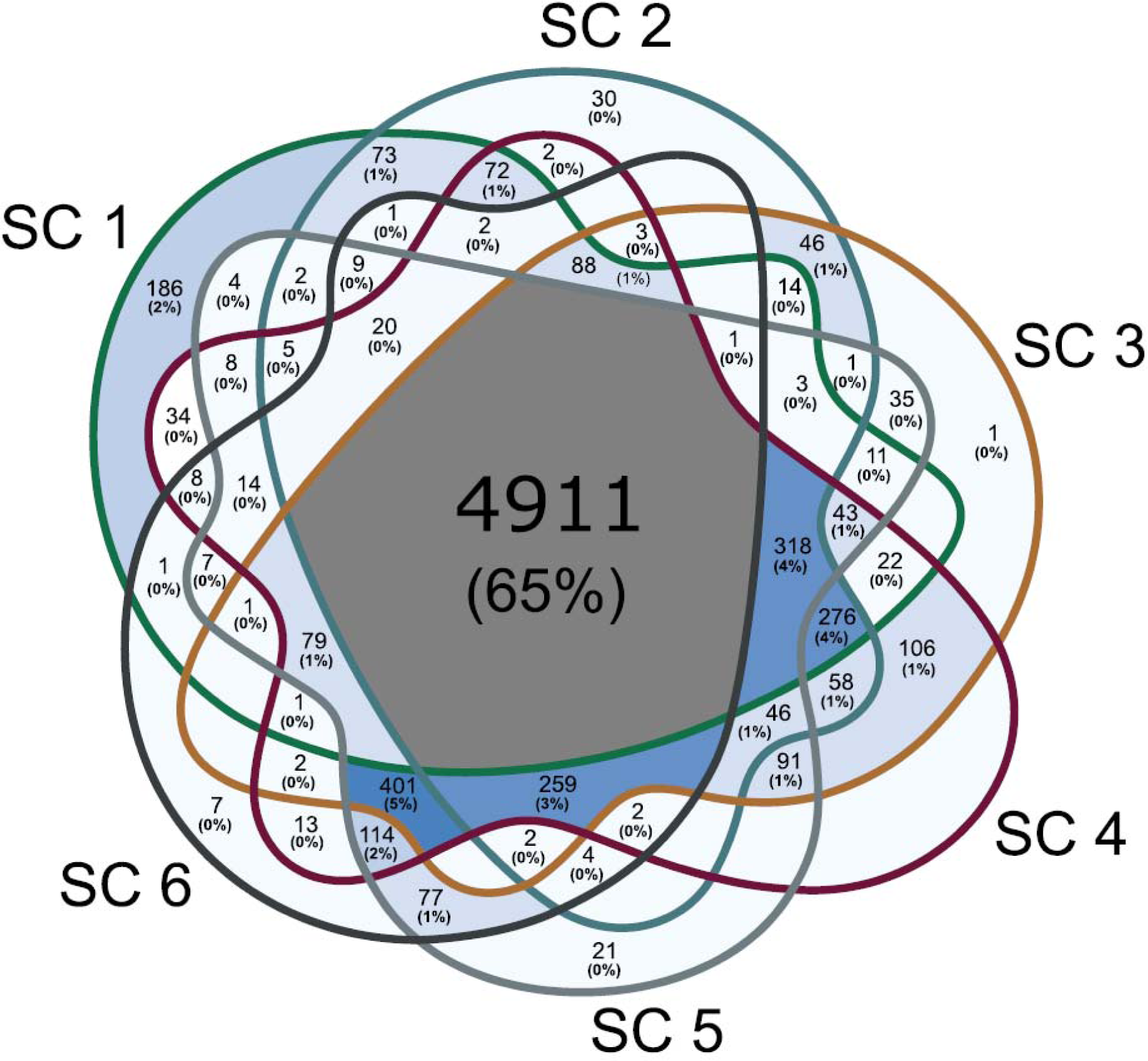
Venn diagram showing unique and shared KOs present in MAGs at different depths.

**Supplementary Fig. 6.**
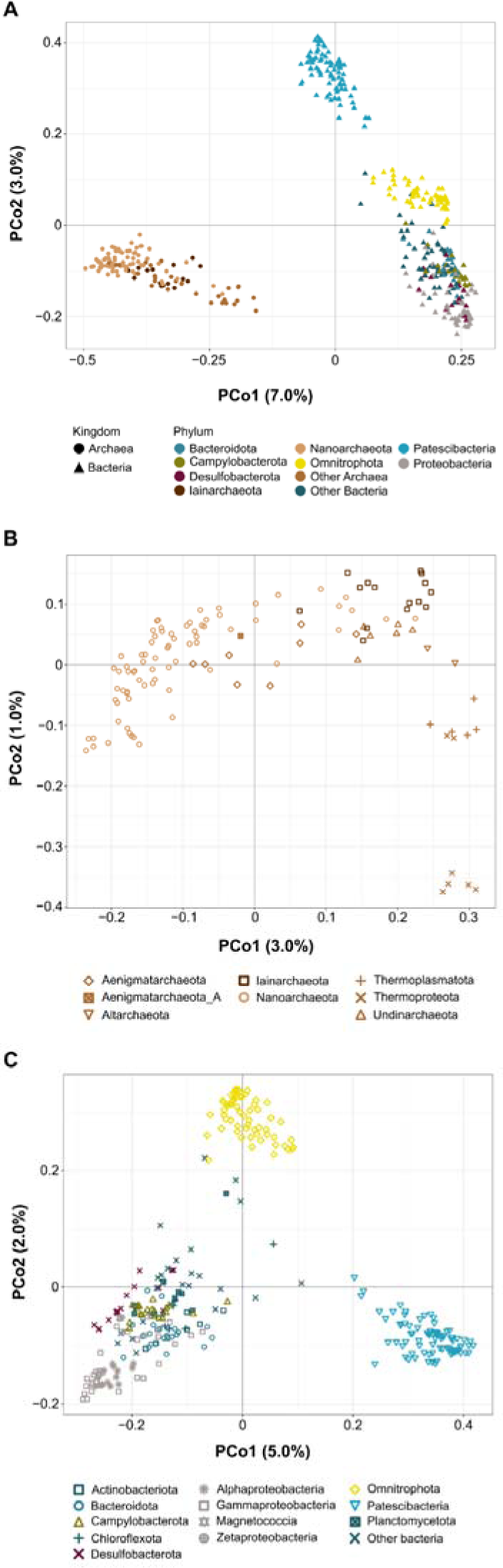
Diversity of MAGs in Sarcophagus cave based on their encoded functional capabilities. **A** whole microbial community, **B** Archaeal community, and **C** Bacterial community.

**Supplementary Fig. 7.**
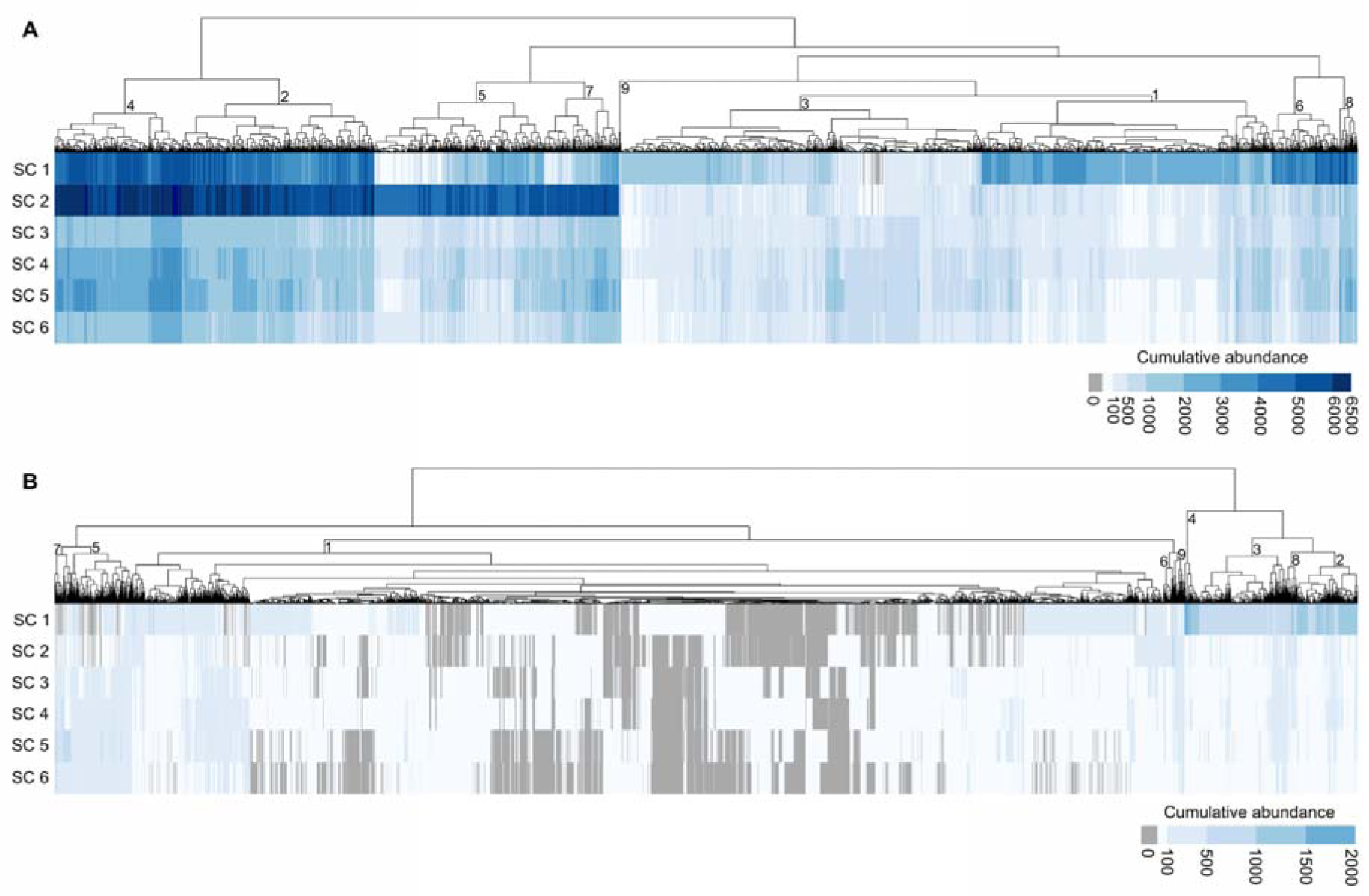
**A** distribution profile of common genes (KOs) **B** rare genes (KOs) in different depths.

**Supplementary Fig. 8.**
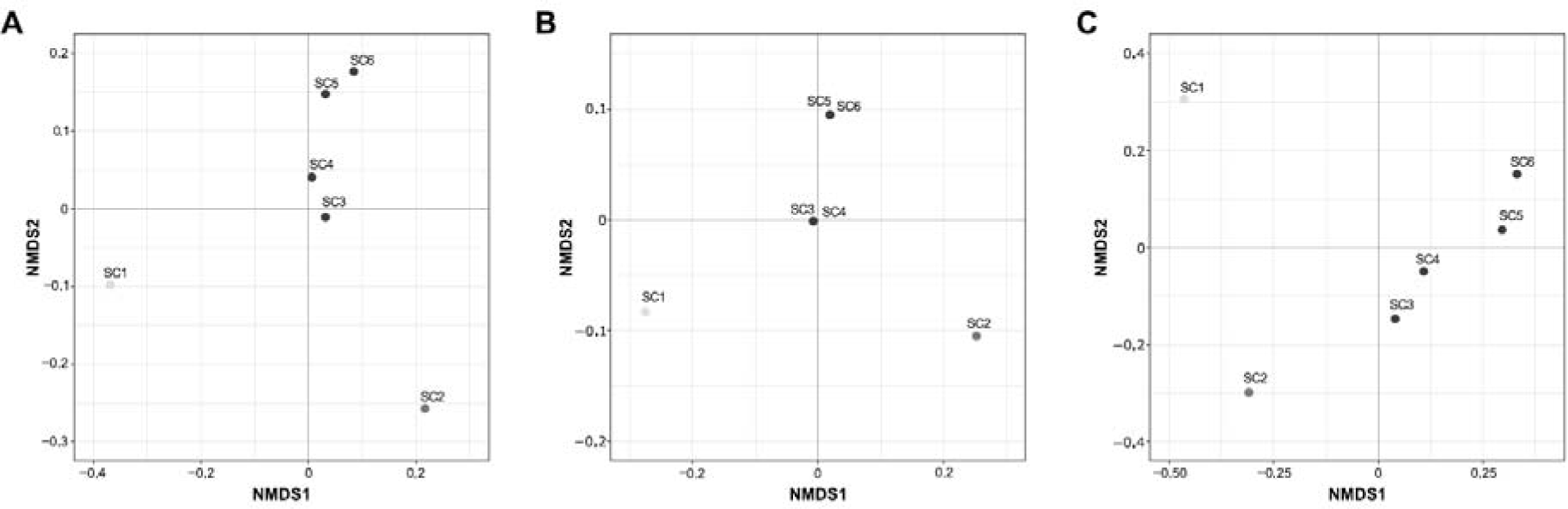
Ordination pattern of KO abundance across different depths. **A** All KOs, **B** common KOs, and **C** rare KOs.

**Supplementary Fig. 9.**
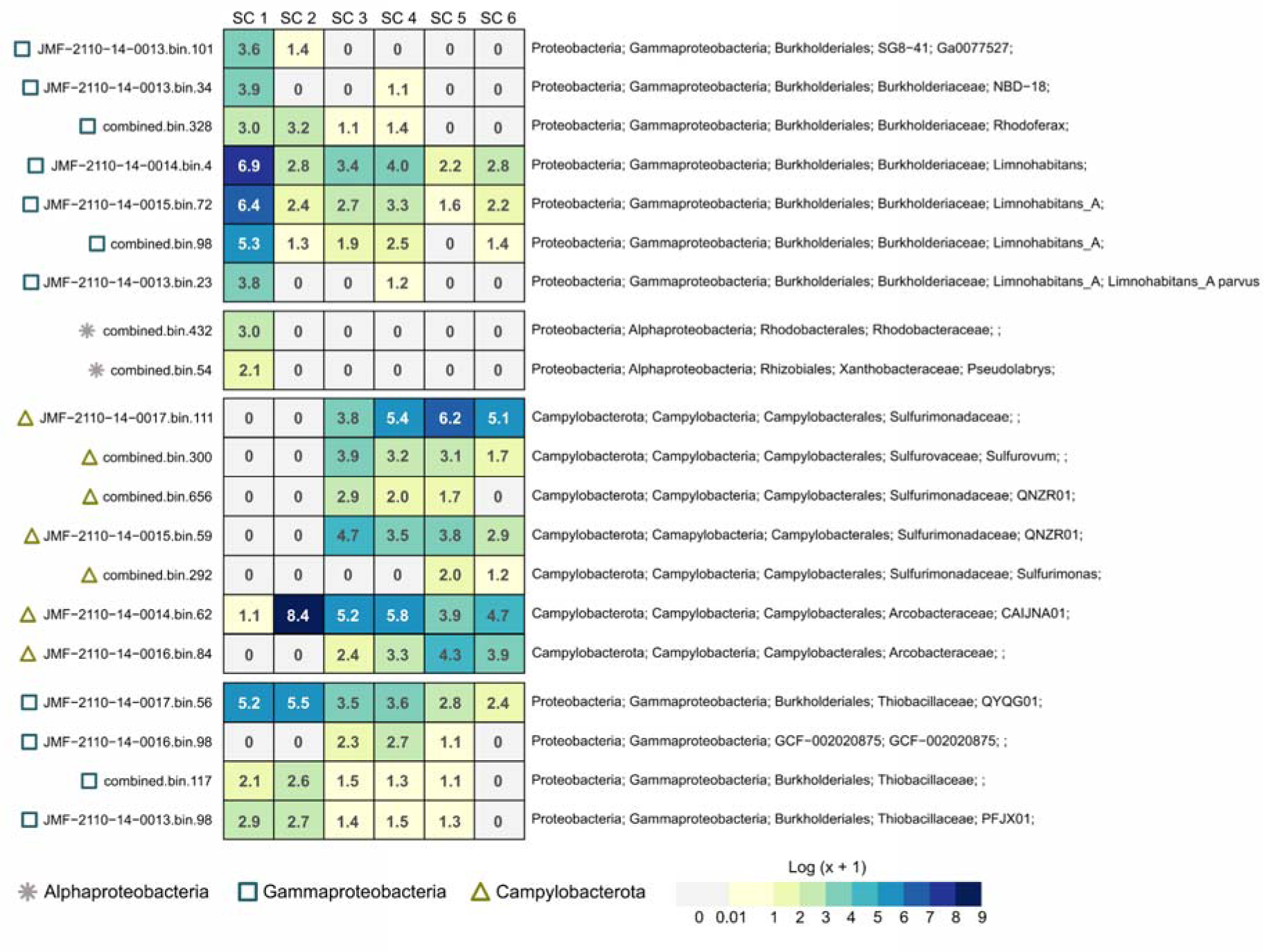
Distribution pattern of MAGs carrying sulfur oxidation (SOX) pathway along the vertical gradient. The abundance of MAGs were log-transformed (log (x+1)). MAGs encoding SOX were separated in four main clusters based on their functional capabilities. The clustering relies on the binary Jaccard distance, quantified using the presence-absence profile of KOs in MAGs.

**Supplementary Fig. 10.**
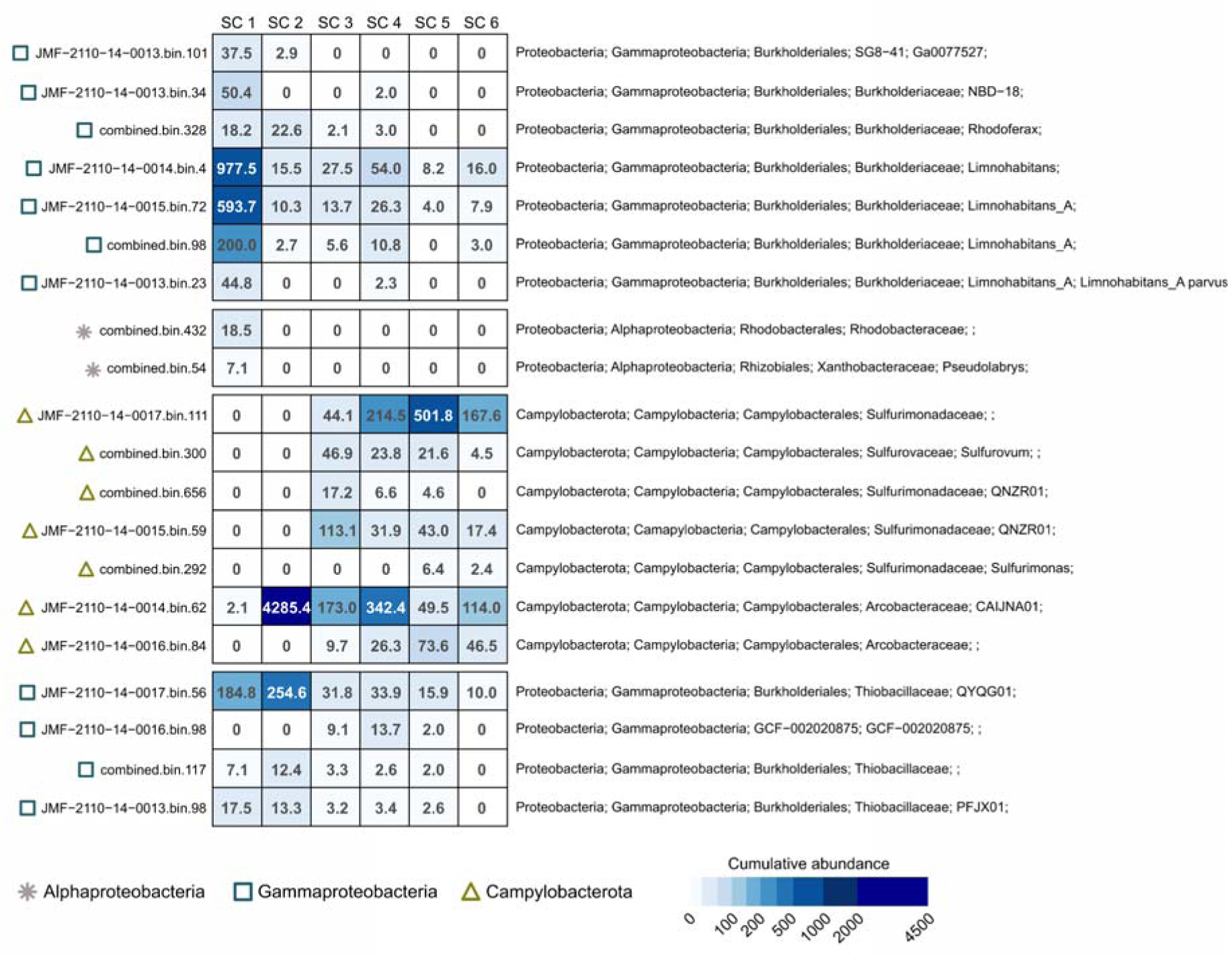
Distribution pattern of MAGs carrying sulfur oxidation (SOX) pathway along the vertical gradient. MAGs encoding SOX were separated in four main clusters based on their functional capabilities. The clustering relies on the binary Jaccard distance, quantified using the presence-absence profile of KOs in MAGs.

**Supplementary Fig. 11.**
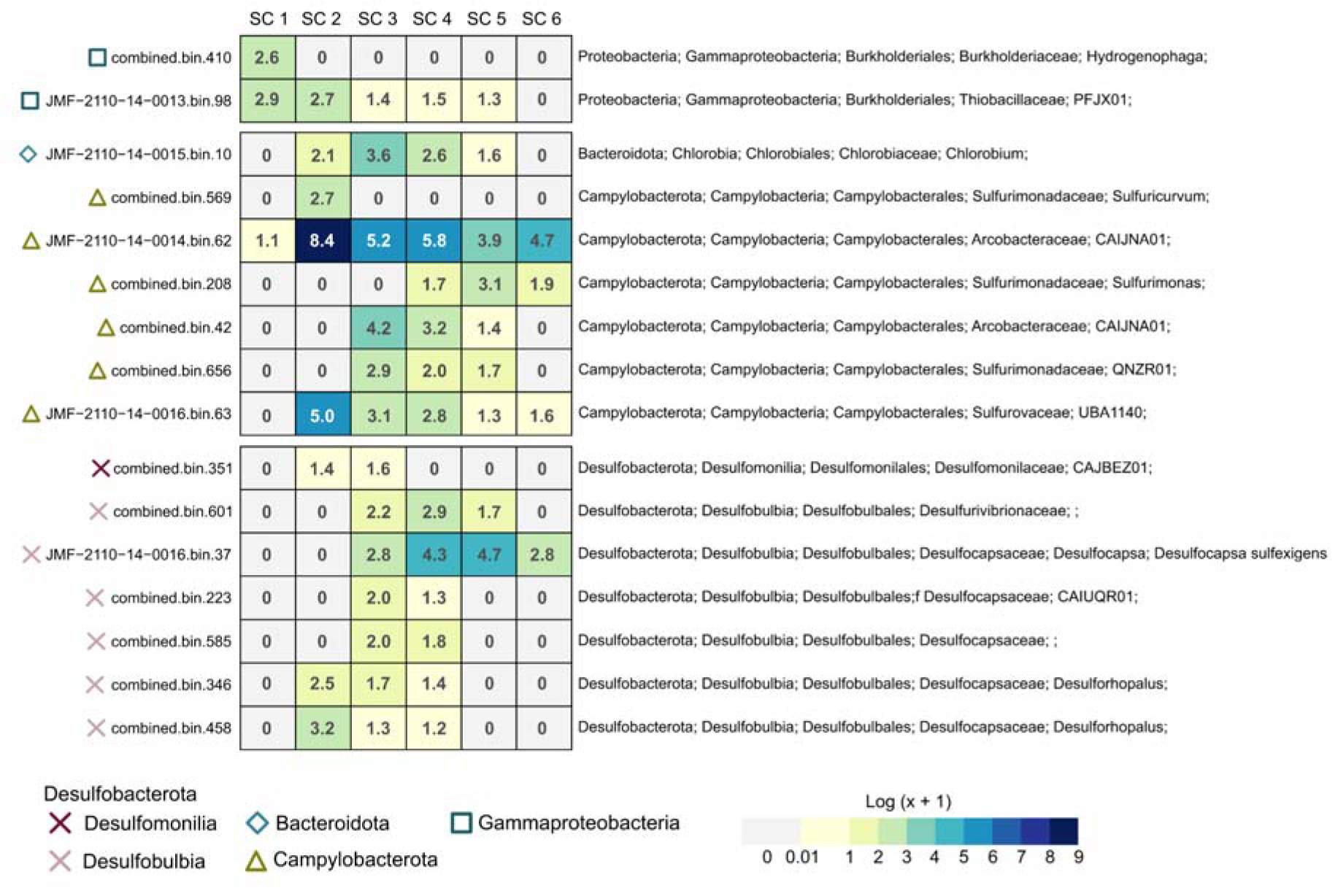
Distribution pattern of MAGs carrying nitrogen fixation (NIF) pathway along the vertical gradient. The abundance of MAGs were log-transformed (log (x+1)). MAGs encoding NIF were separated in three main clusters based on their functional capabilities. The clustering relies on the binary Jaccard distance, quantified using the presence-absence profile of KOs in MAGs.

**Supplementary Fig. 12.**
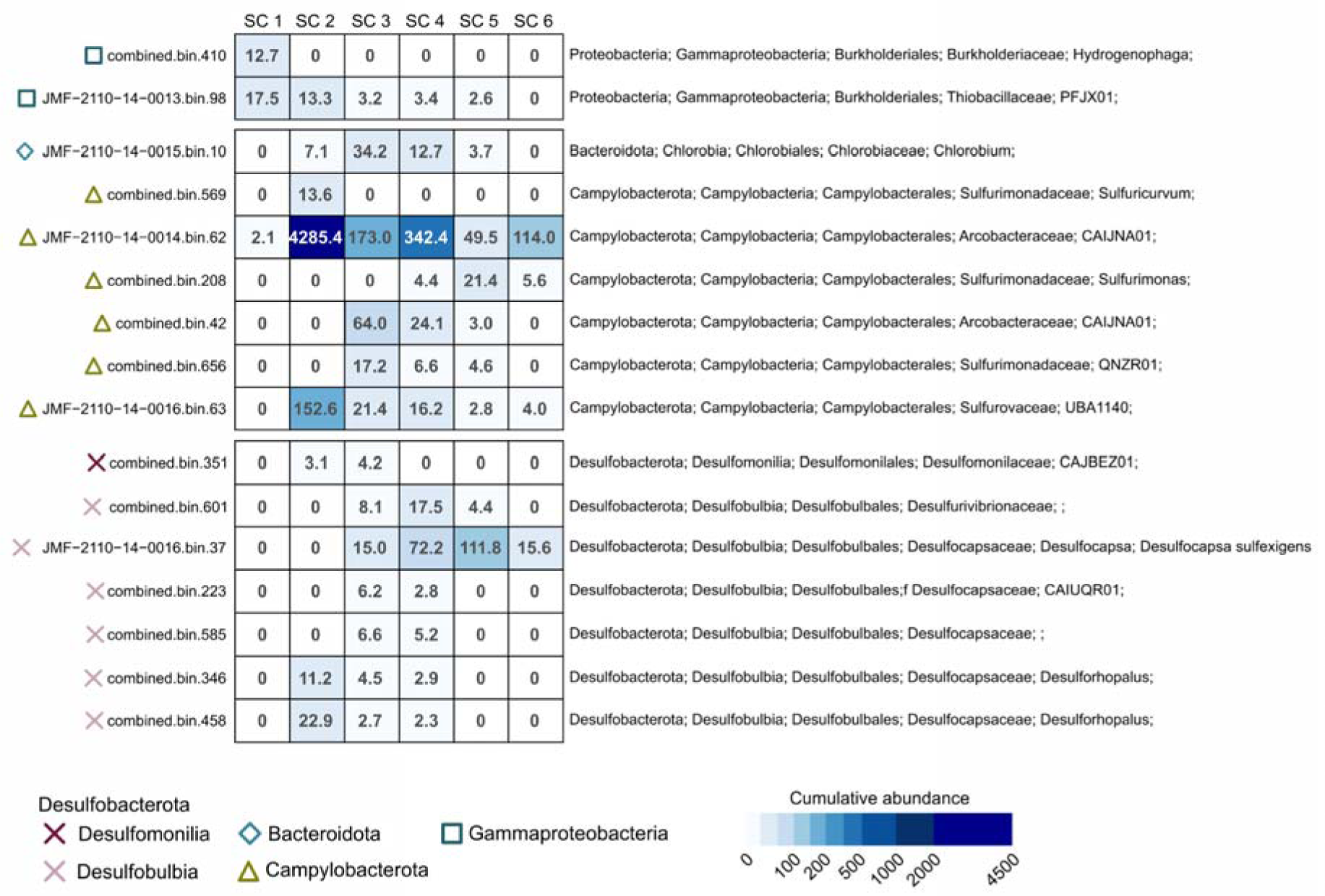
Distribution pattern of MAGs carrying nitrogen fixation (NIF) pathway along the vertical gradient. MAGs encoding NIF were separated in three main clusters based on their functional capabilities. The clustering relies on the binary Jaccard distance, quantified using the presence-absence profile of KOs in MAGs.

**Supplementary Fig. 13.**
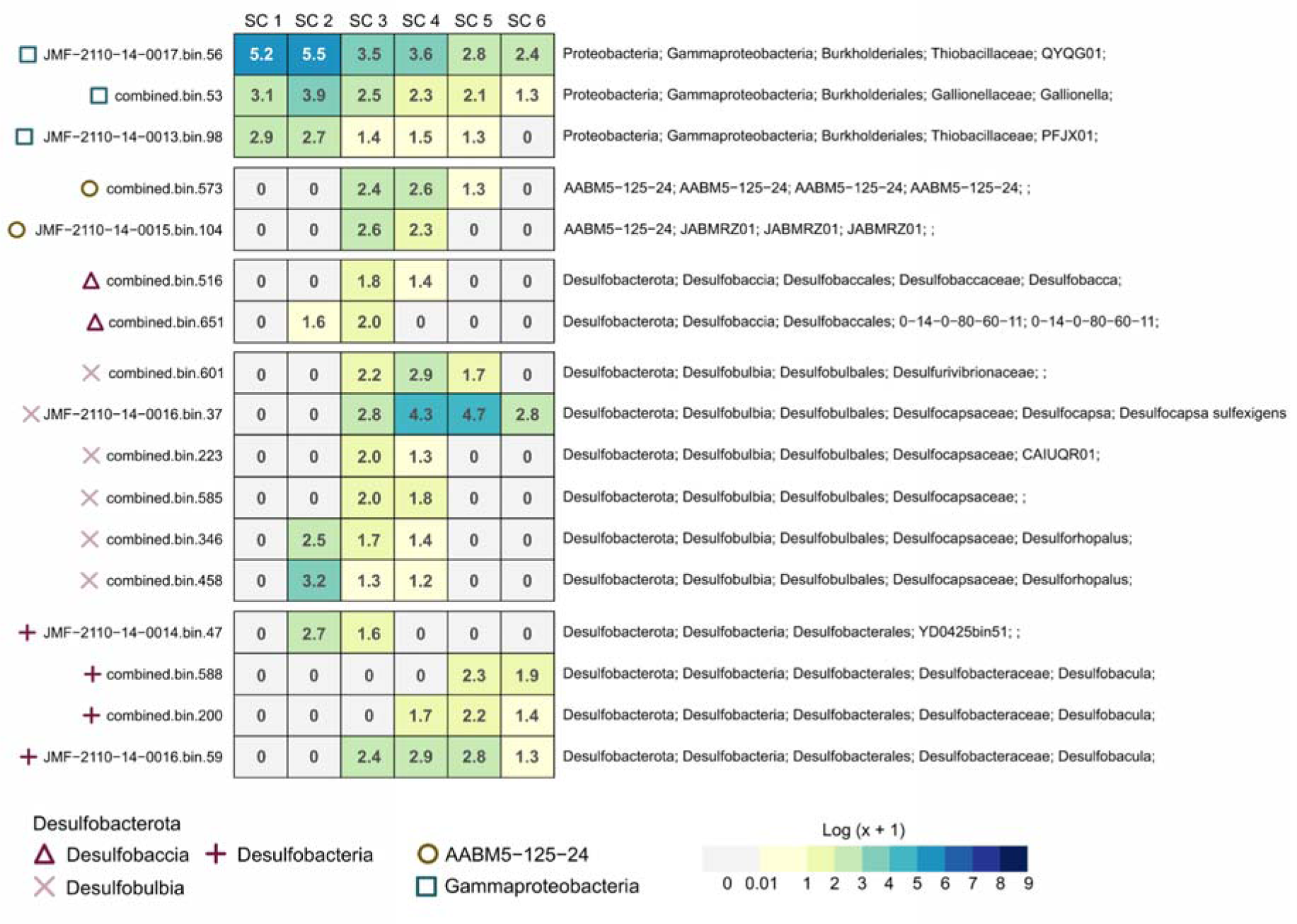
Distribution pattern of MAGs carrying sulfate reduction (DSR) pathway along the vertical gradient. The abundances of MAGs were log-transformed (log (x+1)). MAGs encoding DSR were separated in five main clusters based on their functional capabilities. The clustering relies on the binary Jaccard distance, quantified using the presence-absence profile of KOs in MAGs.

**Supplementary Fig. 14.**
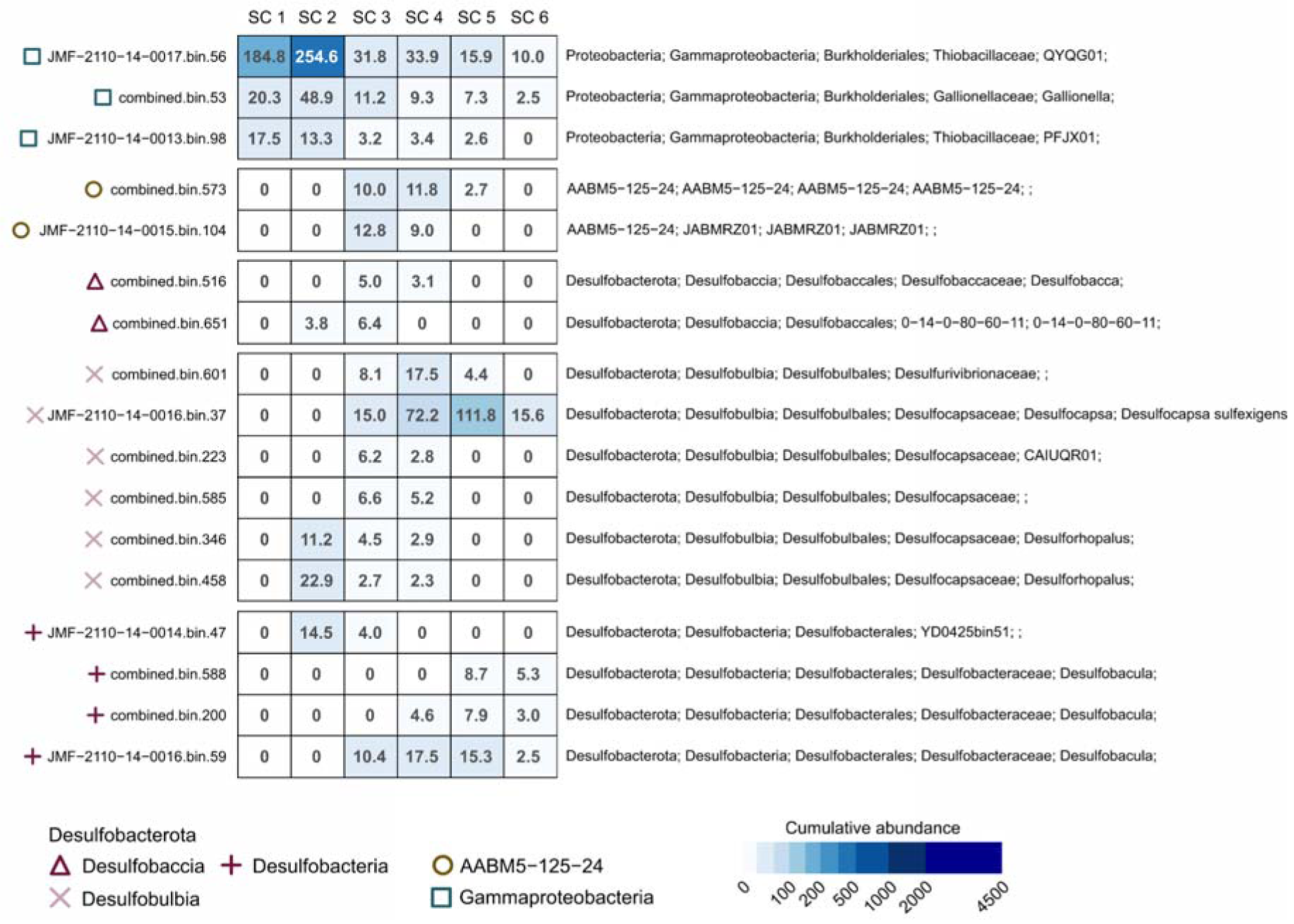
Distribution pattern of MAGs carrying sulfate reduction (DSR) pathway along the vertical gradient. MAGs encoding DSR were separated in five main clusters based on their functional capabilities. The clustering relies on the binary Jaccard distance, quantified using the presence-absence profile of KOs in MAGs.

**Supplementary Fig. 15.**
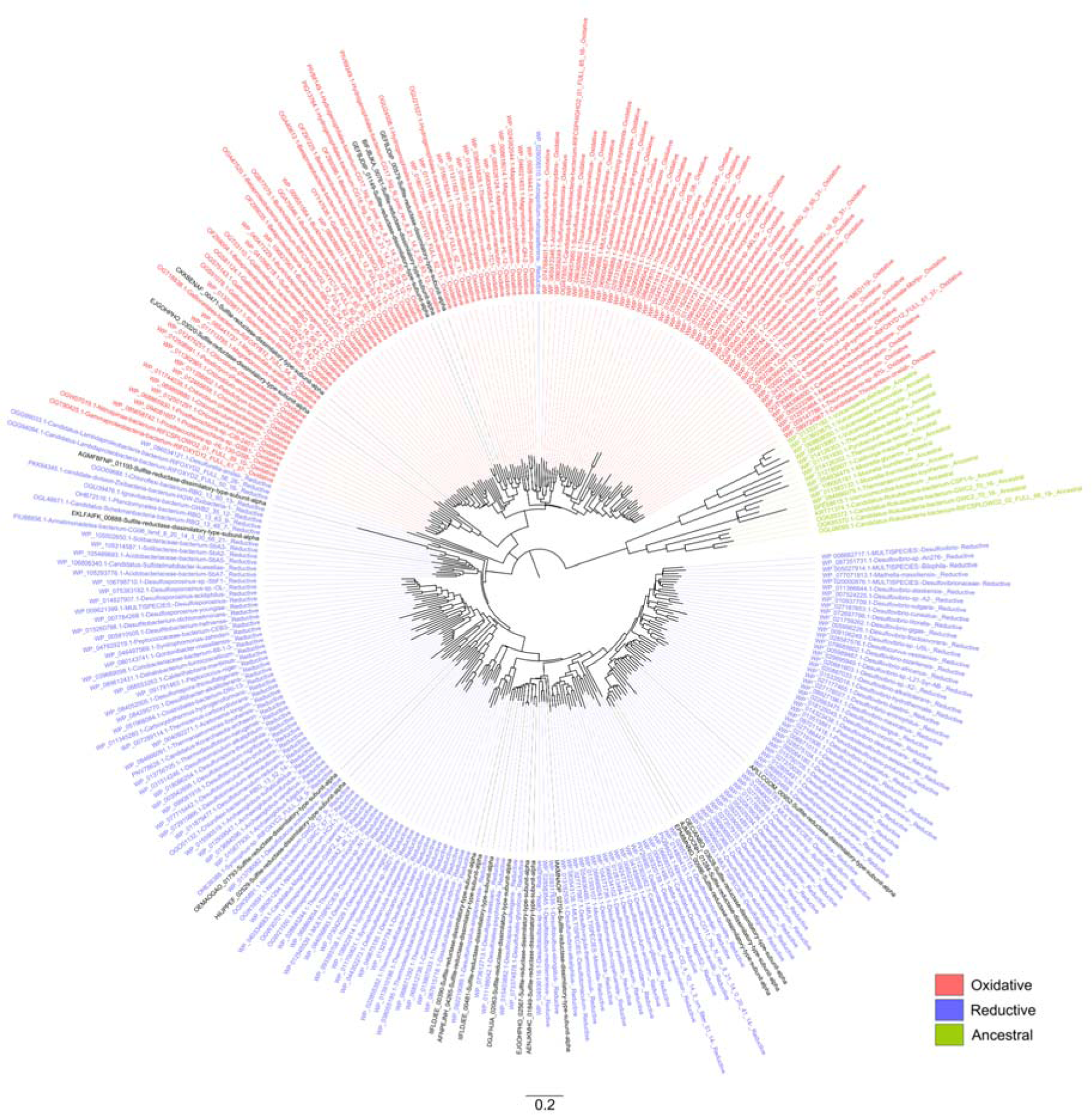
Phylogenetic tree of DSR A genes. Different types of the DSR gene (oxidative, reductive, and ancestral) are colored based on their phylogenetic clustering together with reference sequences.

**Supplementary Fig. 16.**
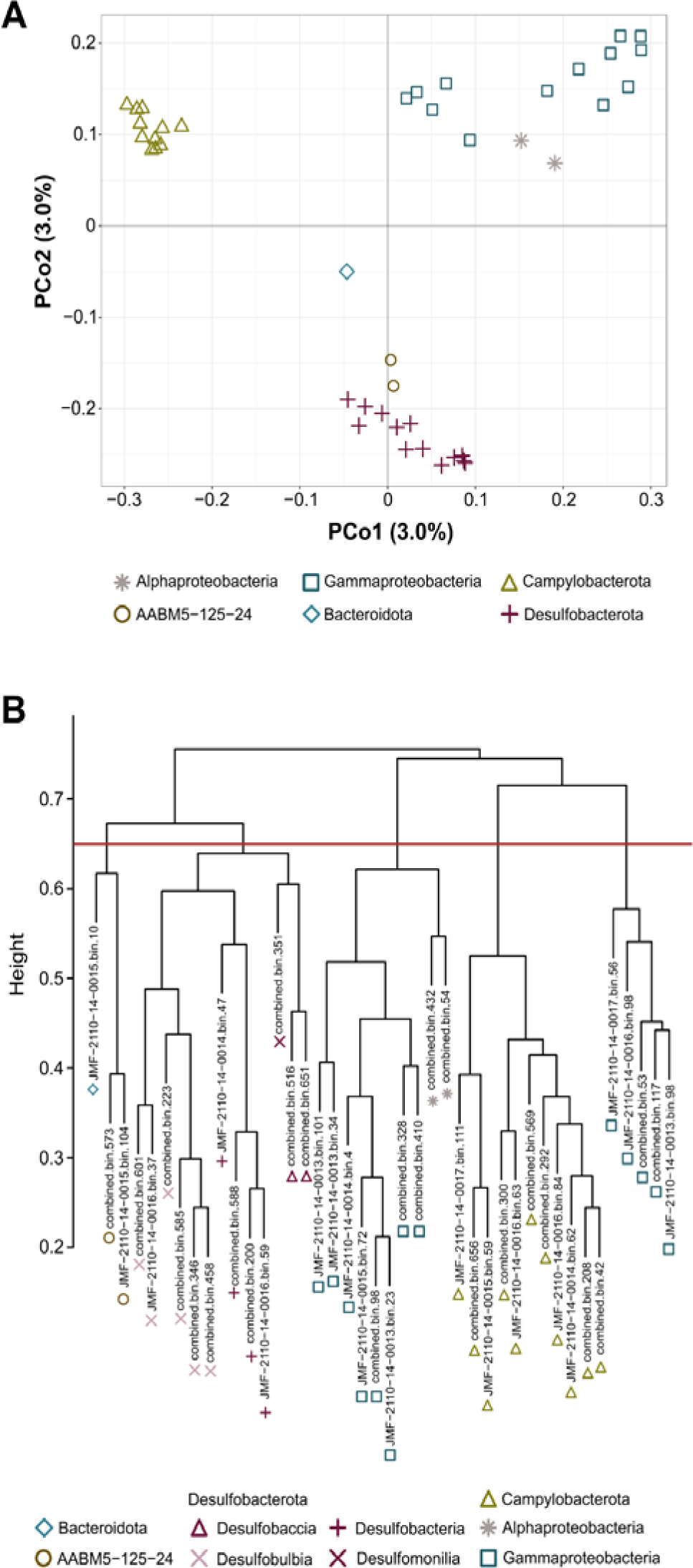
**A** Diversity of functional capabilities encoded in MAGs associated with sulfur oxidation (SOX), sulfate reduction (DSR), and nitrogen fixation (NIF) pathways. **B** Hierarchical relationship between the MAGs and clustering patterns based on their functional capabilities. The red line indicates the operational number of clusters.

**Supplementary Fig. 17.**
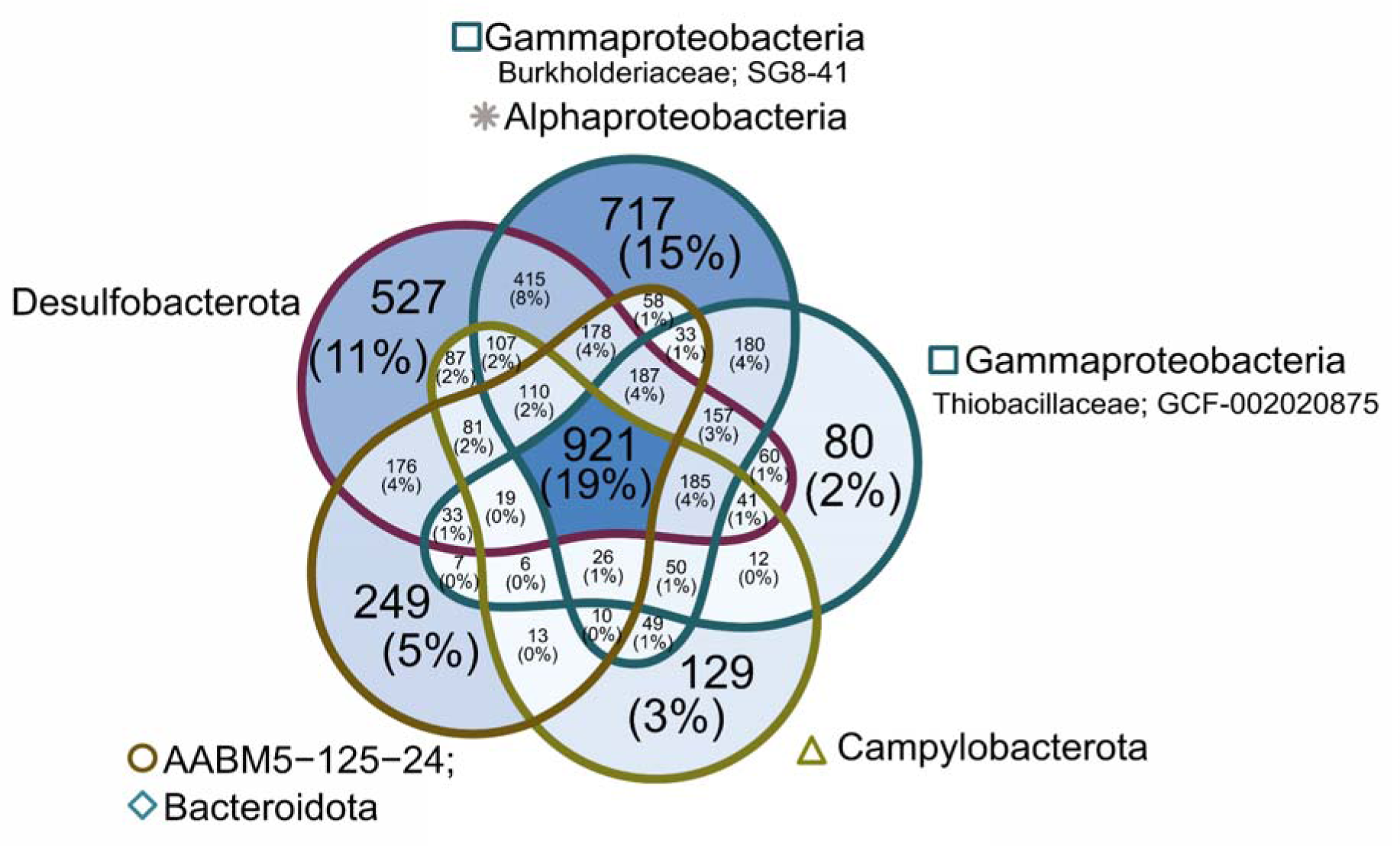
Venn diagram showing unique and shared KOs present in MAGs associated with sulfur oxidation (SOX), sulfate reduction (DSR), and nitrogen fixation (NIF) pathways.

**Supplementary Fig. 18.**
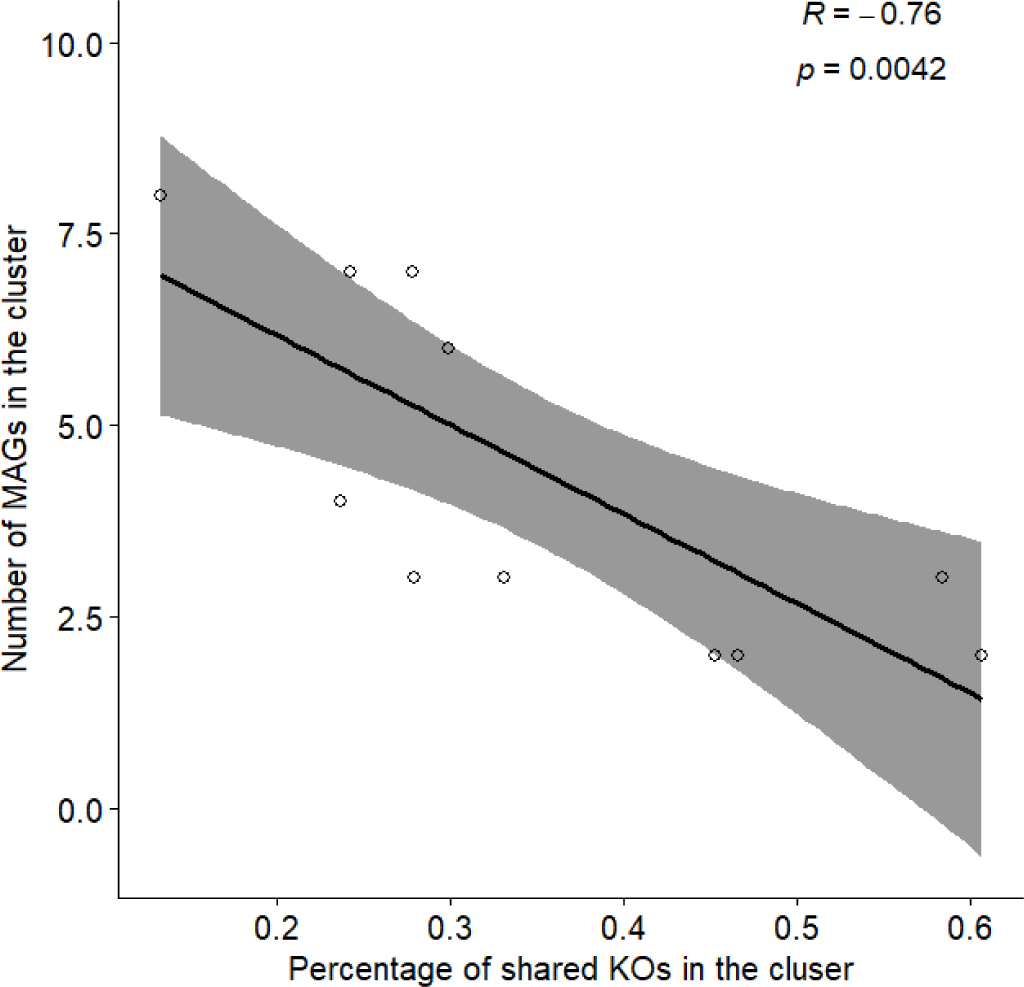
Correlation graph of the percentage of shared KOs and number of MAGs in the cluster.

**Supplementary Fig. 19.**
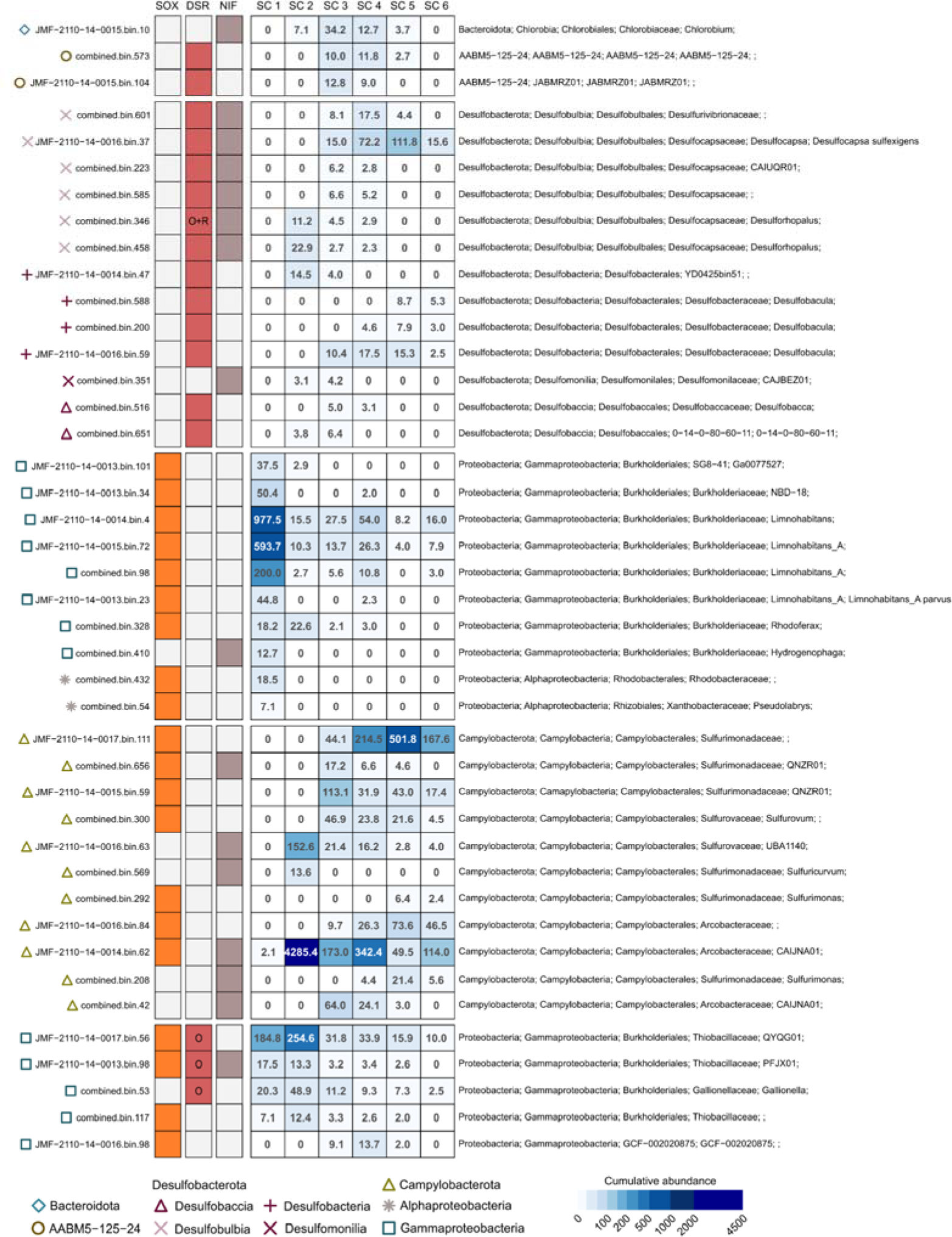
The abundance of MAGs associated with sulfur oxidation (SOX), sulfate reduction (DSR), and nitrogen fixation (NIF) pathways along the vertical gradient. Metabolic analyses of annotated functions separated MAGs containing these functions to five main clusters. The clustering relies on the binary Jaccard distance, quantified using the presence-absence profile of KOs in MAGs. The first three columns indicate the presence of the prominent metabolic module within each MAG.

**Supplementary Fig. 20.**
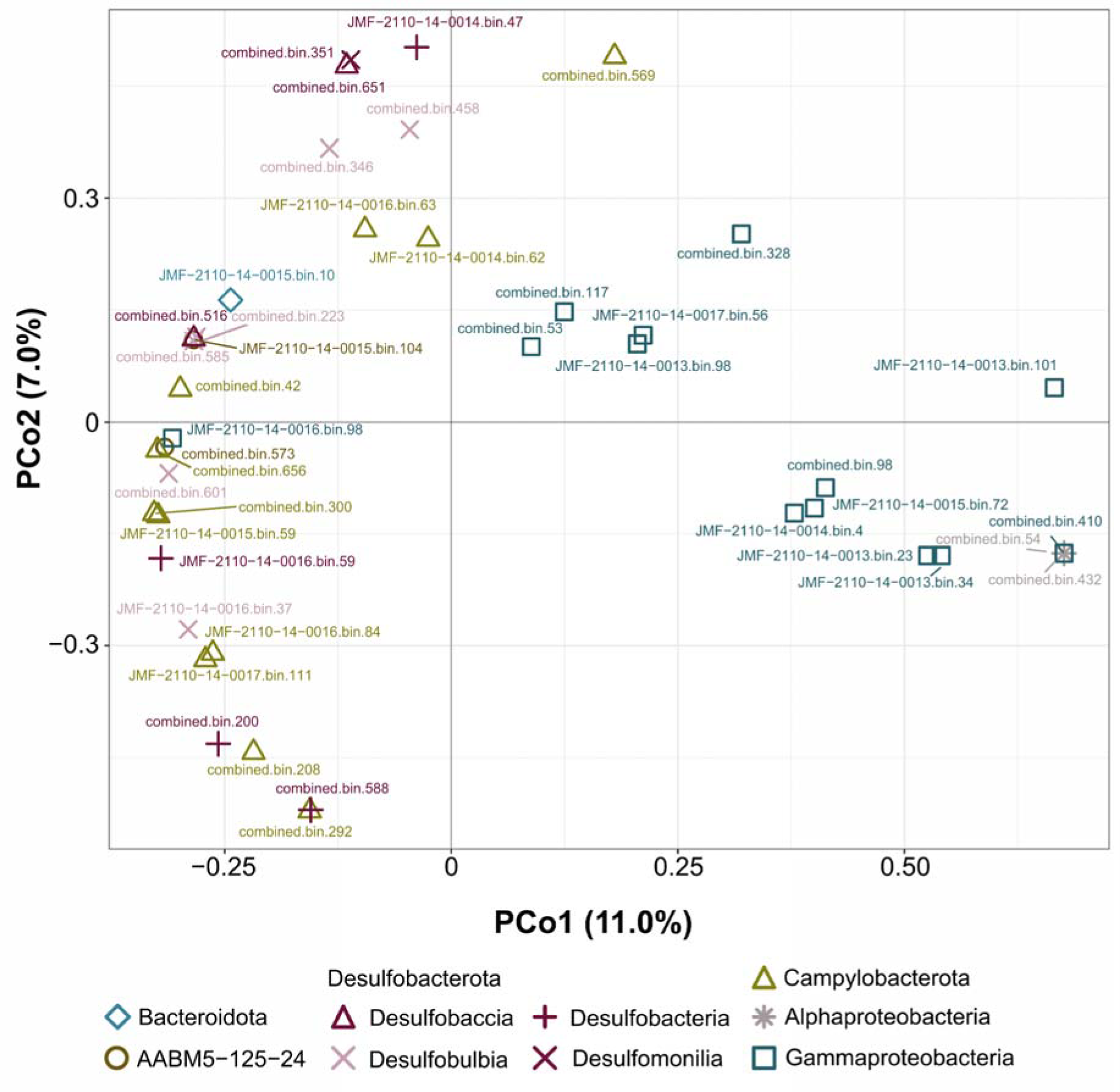
PCoA analyses of the abundance of the MAGs per depth associated with sulfur oxidation (SOX), sulfate reduction (DSR), and nitrogen fixation (NIF) pathways.

## Description of supplementary tables

Supplementary Table 1. Physical and chemical parameters of the studied anchialine cave.

Supplementary Table 2. Sequence abundance, taxonomy, and nucleotide sequences of archaeal and bacterial 16S rRNA gene amplicon sequence variants.

Supplementary Table 3. Relative abundance of most abundant 16S rRNA gene amplicon sequence variants.

Supplementary Table 4. Cumulative abundance, taxonomy and summary statistics of MAGs.

Supplementary Table 5. Abundance-corrected profile of KOs present only in sample SC1.

Supplementary Table 6. Abundance-corrected profile of all KOs.

Supplementary Table 7. Cumulative abundance per depth of selected MAGs with modules (SOX, DSR, NIF).

Supplementary Table 8. Log-transformed abundance per depth of selected MAGs with modules (SOX, DSR, NIF).

Supplementary Table 9. KOs present in MAGs with SOX-related genes.

Supplementary Table 10. KOs present in clusters of MAGs with SOX-related genes. Data shown in Fig. 2 (Venn diagram).

Supplementary Table 11. KOs present in MAGs with NIF-related genes.

Supplementary Table 12. KOs present in clusters of MAGs with NIF-related genes. Data shown in Fig. 2 (Venn diagram).

Supplementary Table 13. KOs present in MAGs with DSR-related genes.

Supplementary Table 14. KOs present in clusters of MAGs with DSR-related genes. Data shown in Fig. 2 (Venn diagram).

Supplementary Table 15. DSR A genes present in selected MAGs with DSR-related genes.

Supplementary Table 16. KOs present in MAGs with SOX-, NIF- and DSR-related genes.

Supplementary Table 17. KOs present in clusters of MAGs with SOX-, NIF- and DSR-related genes. Data shown in Supplementary Fig.16 (Venn diagram).

Supplementary Table 18. Number of shared KOs between MAGs of each cluster.

Supplementary table 19. KOs present in Desulfobacula MAGs with DSR-related genes.

Supplementary Table 20. KOs present in clusters of MAGs presented in Fig.4.

Supplementary Table 21. Description of the metagenomes.

## Supplementary Data

### Supplementary data 1. Correlation of prevalence and cumulative abundance of KOs per depth

For each annotated KO within the MAGs, the cumulative abundance of all MAGs carrying this gene each in depth was assigned as the abundance of that gene in each depth. Prior to further analyses, we have tested whether the prevalence of KOs and their cumulative abundance correlates. We have found high significant positive correlation between prevalence and cumulative abundance of KOs in every depth (Pearson’s correlation, R > 0.8, p < 0.001; **Fig.1&2**).

**Fig. 1.**
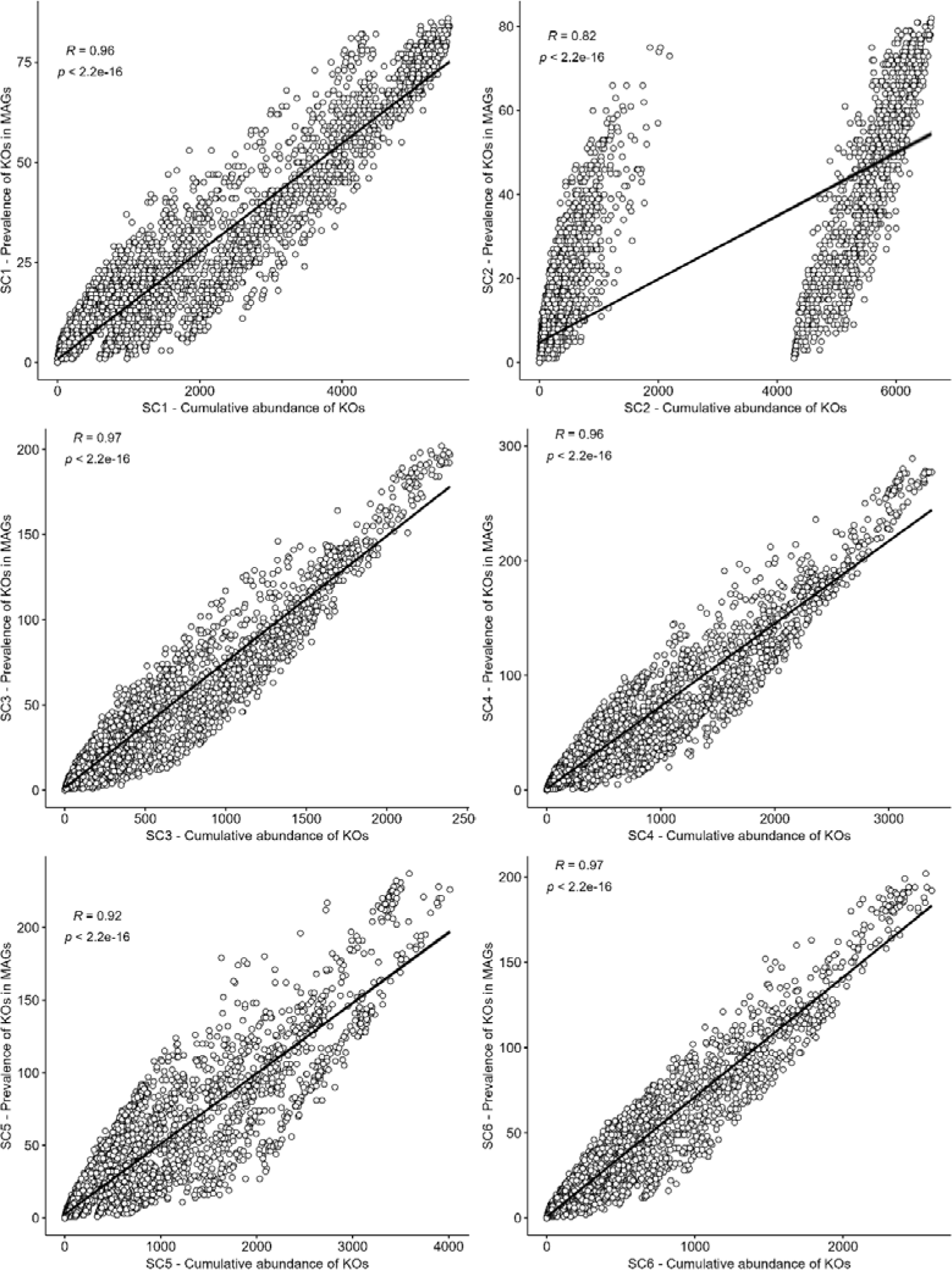
Pearson’s correlation of cumulative abundance of KOs and their prevalence in MAGs per depth.

**Fig. 2.**
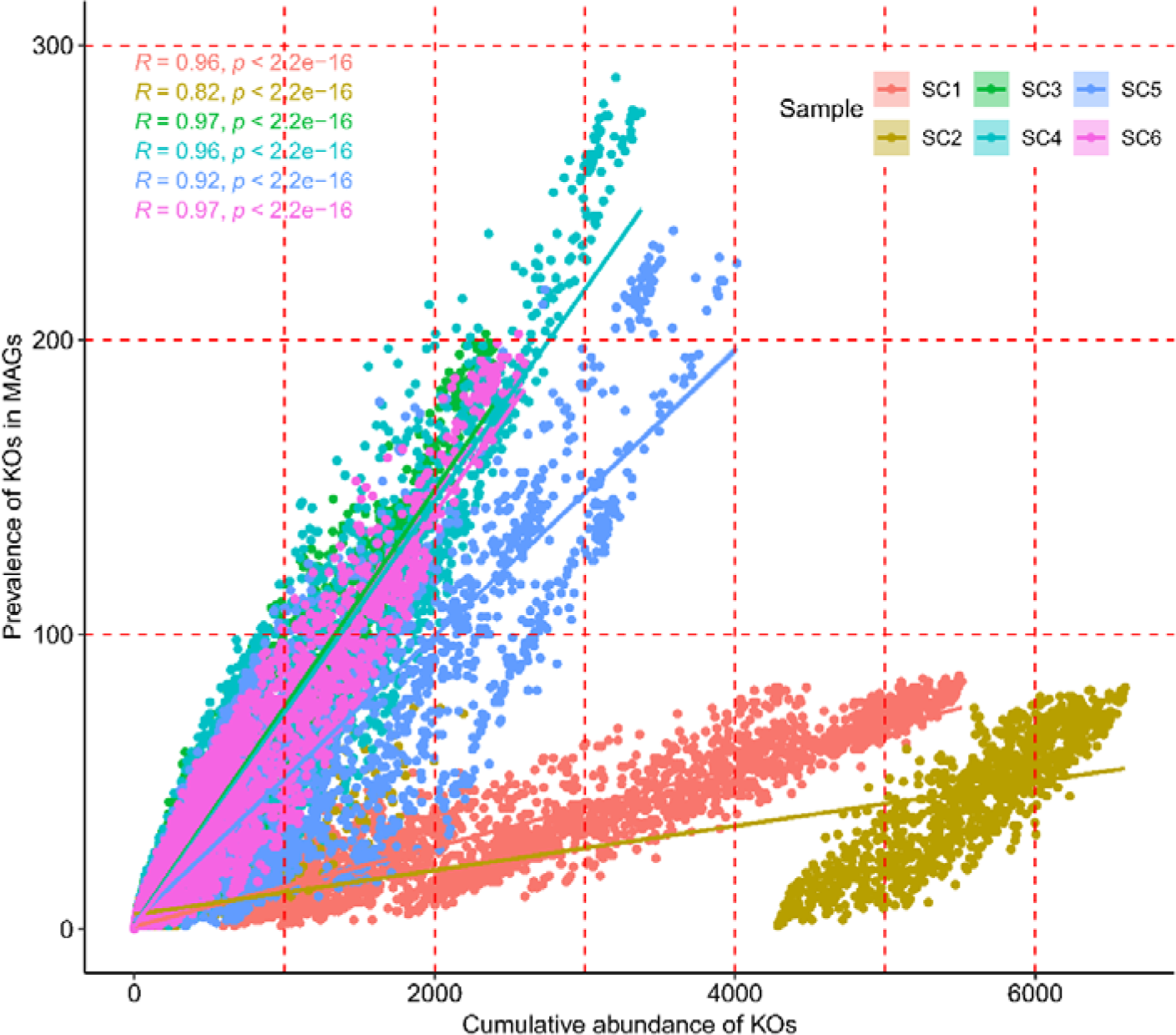
Pearson’s correlation of cumulative abundance of KOs and their prevalence in MAGs per depth.

### Supplementary data 2. Distribution of cumulative abundance of KOs per depth and cutoff point to separate common and rare genes

For each annotated KO within the MAGs, the cumulative abundance of MAGs carrying this gene in each depth was assigned as the abundance of that gene in each depth. The determination of the KOs’ cutoff point involved analyzing the frequency histogram distribution of KOs per depth (**Fig.3**) and in total (**Fig.4**), resulting in a threshold of 2000 based on the summation of their cumulative abundances across all depths. The annotated KOs were further classified into common and rare categories.

**Fig. 3.**
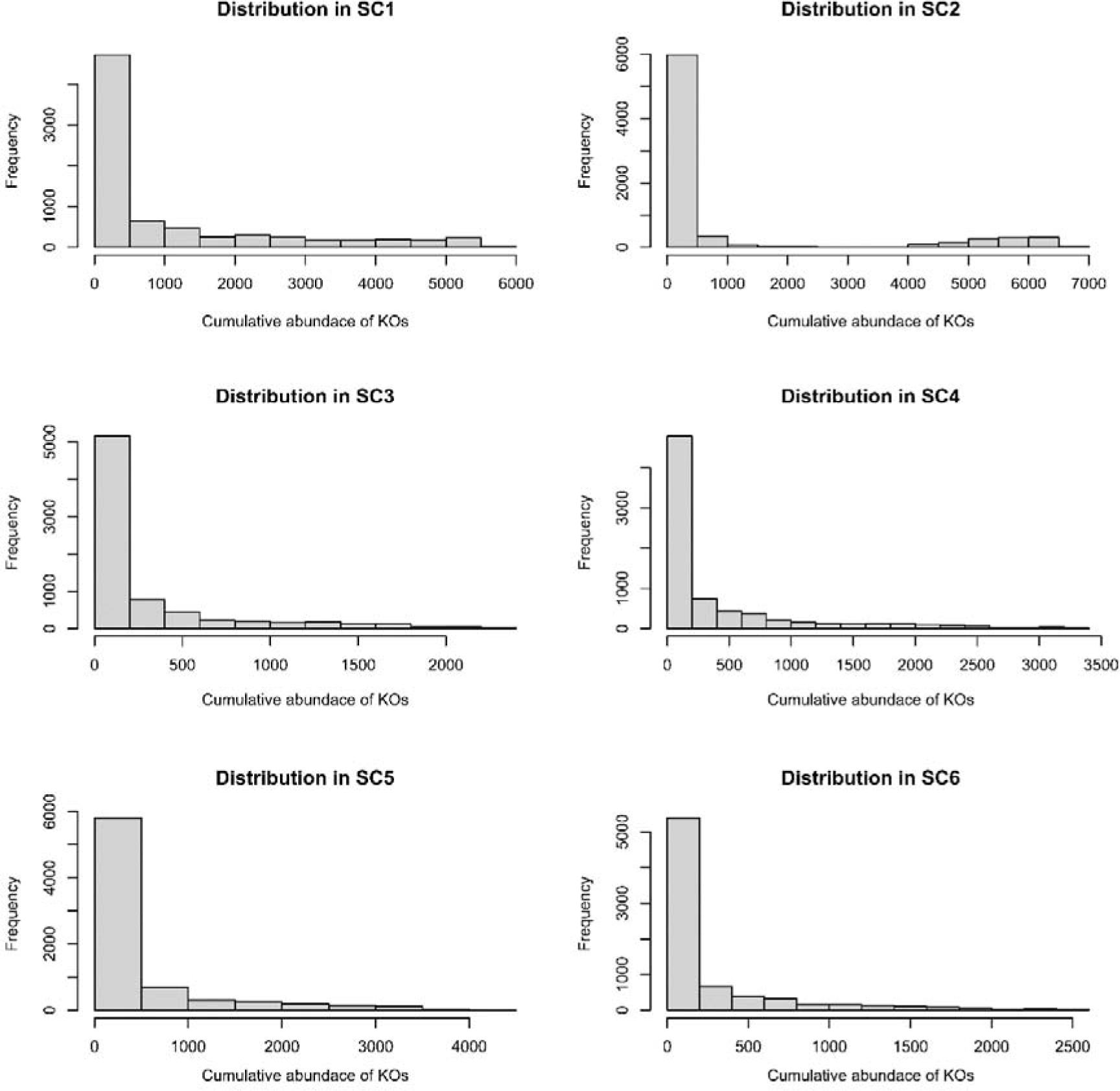
Normal distribution of the cumulative abundance of KOs per depth.

**Fig. 4.**
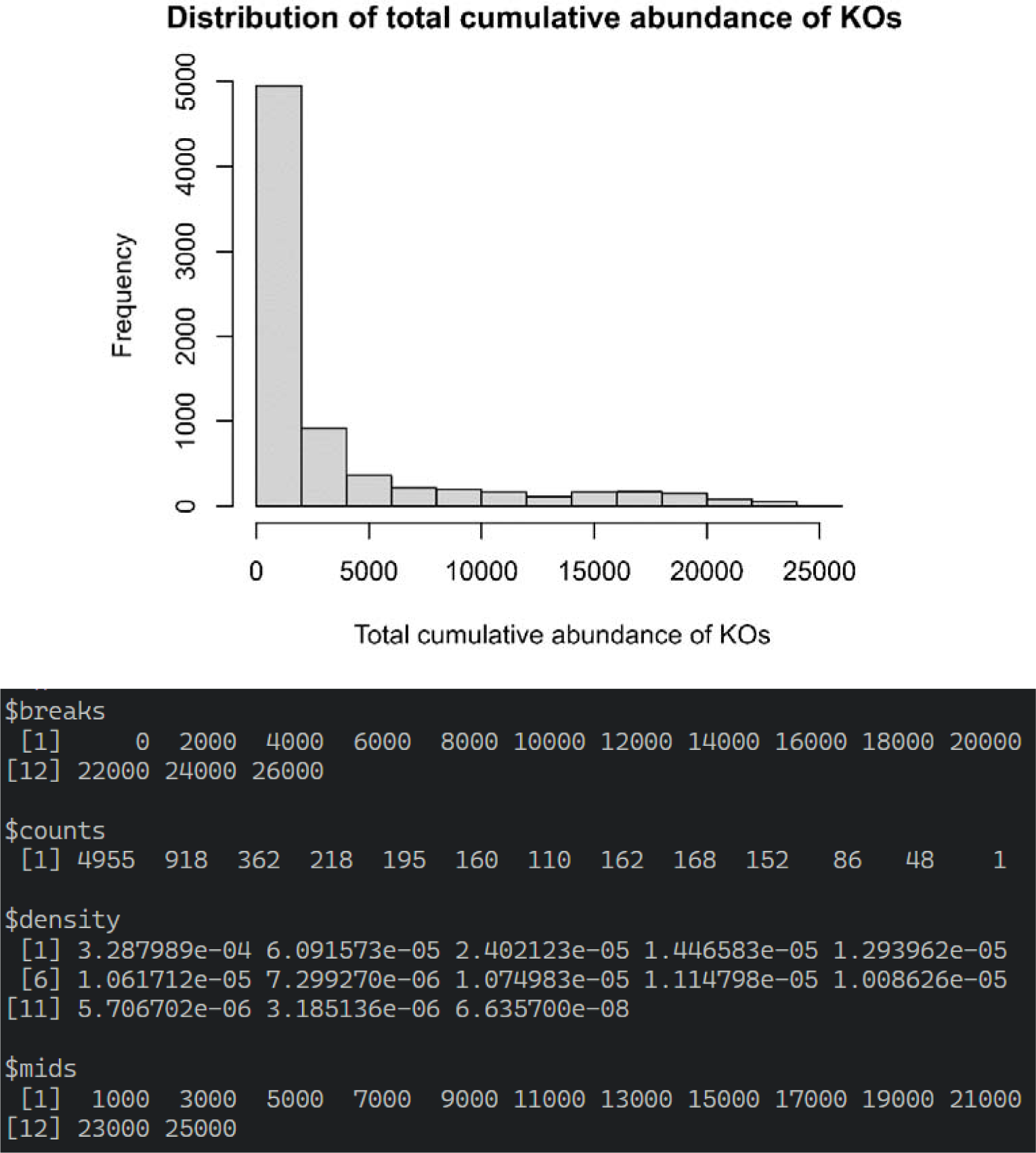
Histogram showing the frequency of total cumulative abundance of KOs and summary. Based on these results we made a cutoff at 2000.

### Supplementary data 3. Diversity pattern of KO abundance across deeper strata

In Supplementary Fig.8, we have shown that the ordination of abundance-corrected metabolic profiles of each depth based on their dissimilarity for all KOs is mainly driven by the differential abundance of MAGs at each depth. While the deeper strata samples have rather overlapped and separated from SC1 and SC2, we further tested their ordination based on all KOs, common and rare KOs (**Fig. 5**). Highest dissimilarity of deeper strata was between the communities based on the rare KOs, specifically between the SC5 and SC6 samples.

**Fig. 5.**
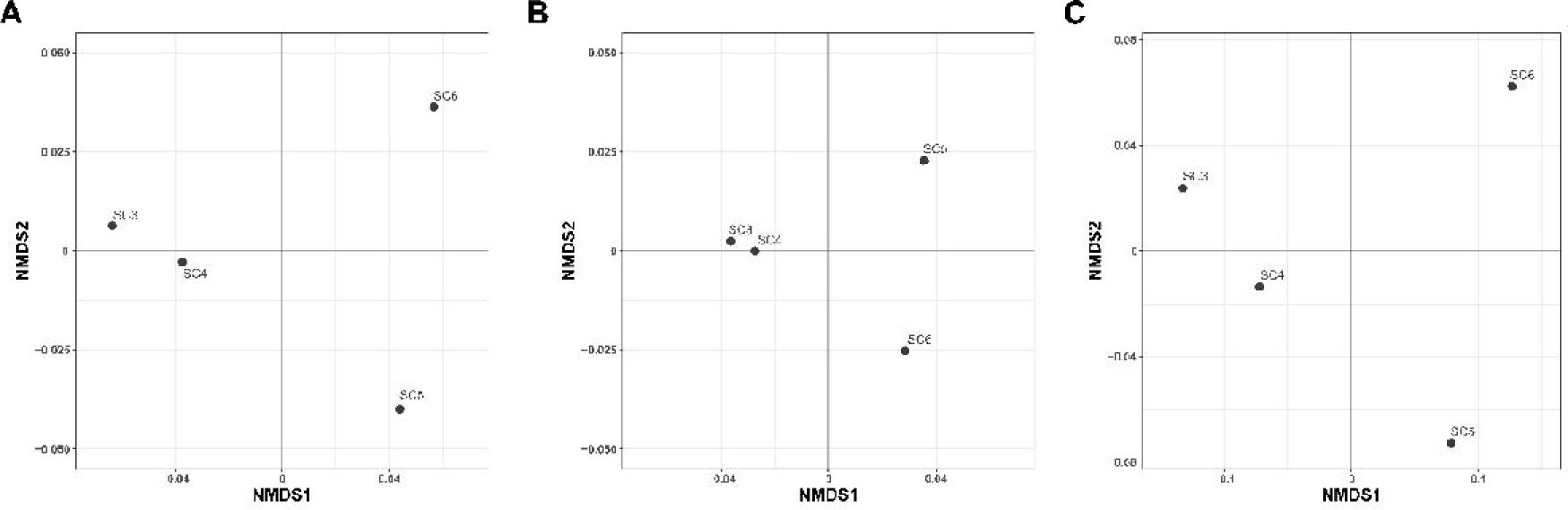
Diversity pattern of KO abundance across deeper strata. **A** All KOs, **B** common KOs, and **C** rare KOs.

### Supplementary data 4. Clustering of MAGs based on the presence/absence of KOs

Results of PERMANOVA per module (SOX, DSR and NIF) and all modules (42 MAGs) confirm the results of the hierarchical clustering method shown in Fig.2 and Supplementary Fig.16 (permutations = 999; **Table 1**). Jaccard distance was calculated based on the presence/absence of KOs per MAG. Based on the dataset of KOs within the SOX, DSR, NIF and all modules.

**Table 1.**
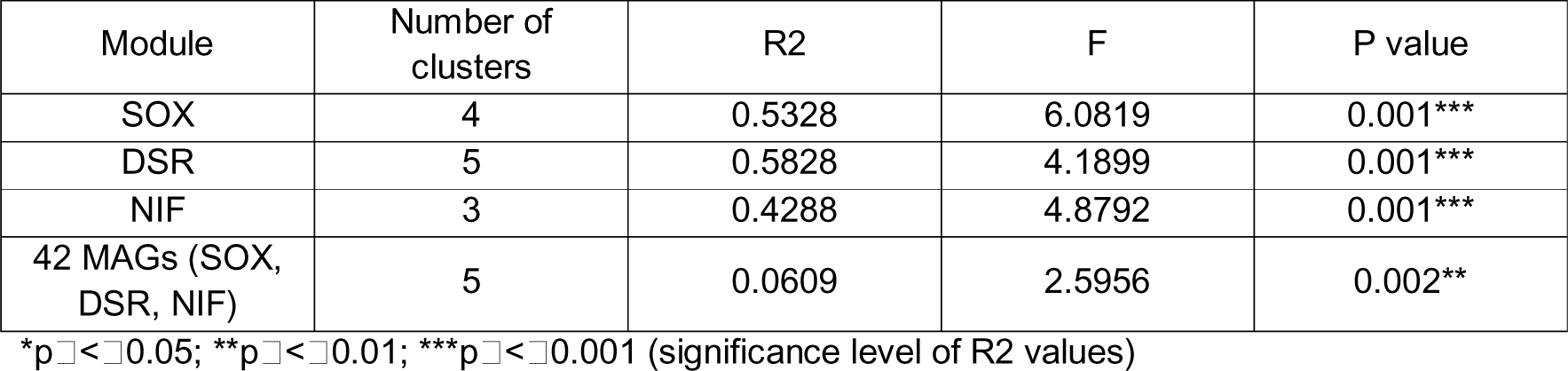
Results of the PERMANOVA analysis on the presence/absence of KOs in MAGs (Jaccard distance matrix) and clusters (shown in dendrograms Fig 2. and Supplementary Fig.16).

| Module | Number of clusters | R2 | F | P value |
| --- | --- | --- | --- | --- |
| SOX | 4 | 0.5328 | 6.0819 | 0.001*** |
| DSR | 5 | 0.5828 | 4.1899 | 0.001*** |
| NIF | 3 | 0.4288 | 4.8792 | 0.001*** |
| 42 MAGs (SOX, DSR, NIF) | 5 | 0.0609 | 2.5956 | 0.002** |

\* $p \leq 0.05$ ; \*\* $p \leq 0.01$ ; \*\*\* $p \leq 0.001$ (significance level of R2 values)

### Supplementary data 5. Correlation of annotated metabolic profile and relative abundance of MAGs to their taxonomic affiliation within studied modules (SOX, DSR, NIF and all 42 MAGs)

The clustering of MAGs based on the presence/absence of KOs resulted in clusters aligned to their taxonomic affiliation. To further test whether the relative abundance of MAGs per depth aligns to their taxonomic affiliation, and thus the KO based clustering, we have applied two methods: PERMANOVA and Mantel test. Considering only MAGs with studied modules (SOX, DSR, NIF), we have calculated the relative abundance of MAGs per depth and their dissimilarity based on the Bray-Curtis distance (**Fig. 6**). First, we tested with PERMANOVA whether their relative abundance aligns with their taxonomic affiliation (shown in dendrograms Fig 2. and Supplementary Fig. 16). Relative abundance of MAGs showed low but significant grouping based on their taxonomic affiliation within all studied modules (**Table 2**). Further, we have tested whether the metabolic profile of MAGs correlates with their relative abundance with the Mantel test and corr.test (package psych). The highest significant correlation was found between the metabolic and abundance profile of SOX-carrying MAGs (**Table 3**).

**Fig. 6.**
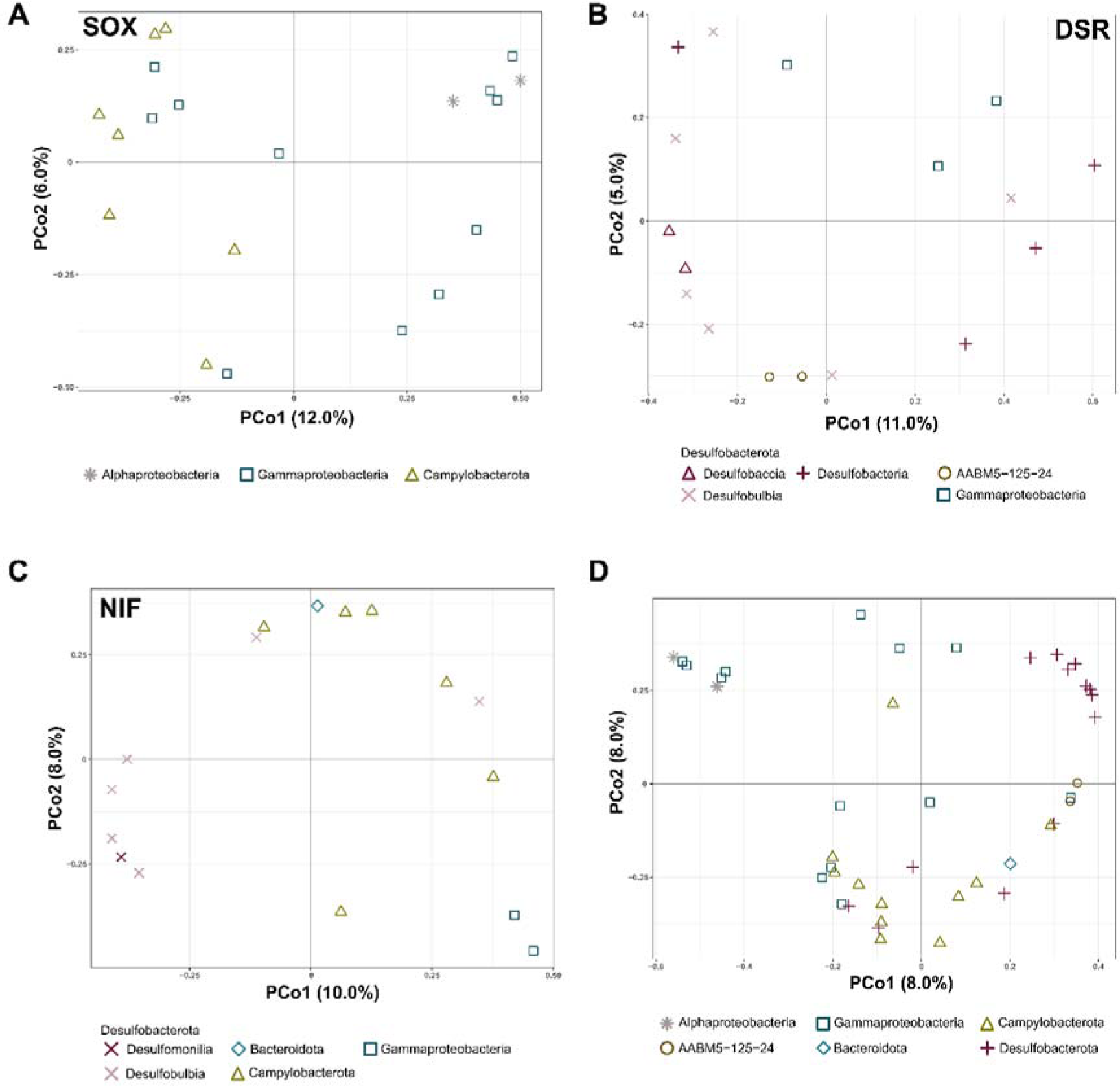
Diversity of relative abundance of MAGs associated with **A** sulfur oxidation (SOX), **B** sulfate reduction (DSR), **C** nitrogen fixation (NIF) and **D** all studied pathways.

**Table 2.**
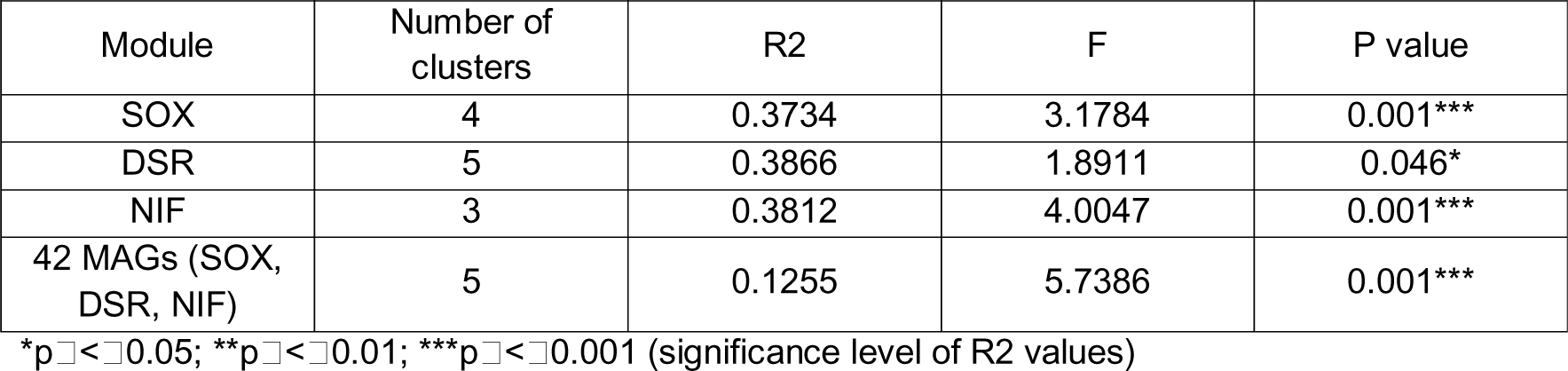
Results of the PERMANOVA analysis of the matrix of relative abundance (1/100%) of MAGs per depth (Bray-Curtis dissimilarity) and clusters (shown in dendrograms Fig 2 and Supplementary Fig. 16).

| Module | Number of clusters | R2 | F | P value |
| --- | --- | --- | --- | --- |
| SOX | 4 | 0.3734 | 3.1784 | 0.001*** |
| DSR | 5 | 0.3866 | 1.8911 | 0.046* |
| NIF | 3 | 0.3812 | 4.0047 | 0.001*** |
| 42 MAGs (SOX, DSR, NIF) | 5 | 0.1255 | 5.7386 | 0.001*** |

\* $p < 0.05$ ; \*\* $p < 0.01$ ; \*\*\* $p < 0.001$ (significance level of R2 values)

**Table 3.** Results of the Mantel test and corr.test (package psych) following the correlation of the annotated metabolic profile (presence/absence of KOs per MAG (Jaccard distance)) and relative abundance of MAGs (relative abundance (1/100%) of MAGs per depth (Bray-Curtis distance)).

| Module | Mantel test<br>(Spearman's rank) |  | Corr.test |  |
| --- | --- | --- | --- | --- |
|  | R value | P value | R value | P value |
| SOX | 0.4545 | 0.001*** | 0.3766 | 0.001*** |
| DSR | 0.09528 | 0.218 | 0.1699 | 0.048* |
| NIF | 0.2977 | 0.026* | 0.245 | 0.007** |
| 42 MAGs (SOX, DSR, NIF) | 0.3515 | 0.001*** | 0.352 | 0.001*** |

\* $p < 0.05$ ; \*\* $p < 0.01$ ; \*\*\* $p < 0.001$ (significance level of R values)

### Supplementary data 6. Influence of incompleteness of MAGs on their metabolic profile

Prior to the analyses based on the chosen metabolic modules (SOX, DSR and NIF), to test if incompleteness of MAGs affects the analyses based on the presence/absence of KOs within the MAGs with SOX, DSR and NIF module, Mantel test was performed based on Euclidean (**Table 4**) and Jaccard (Table 5) distance between raw and subsampled dataset using Spearman’s rank correlation.

**Table 4.** Results of the Mantel test performed between the Euclidean distance matrices of the raw and subsampled datasets of KOs.

| Module | Distance 1 | Distance 2 | Mantel correlation<br>r | P value |
| --- | --- | --- | --- | --- |
| SOX | Raw KO dataset | Subsampled KO dataset | 0.897 | 0.001*** |
| DSR | Raw KO dataset | Subsampled KO dataset | 0.866 | 0.001*** |
| NIF | Raw KO dataset | Subsampled KO dataset | 0.968 | 0.001*** |

\* $p < 0.05$ ; \*\* $p < 0.01$ ; \*\*\* $p < 0.001$ (significance level of r values)

**Table 5.** Results of the Mantel test performed between the Jaccard distance matrices of raw and subsampled dataset of KOs.

| Module | Distance 1 | Distance 2 | Mantel correlation<br>r | P value |
| --- | --- | --- | --- | --- |
| SOX | Raw KO dataset | Subsampled KO dataset | 0.499 | 0.001*** |
| DSR | Raw KO dataset | Subsampled KO dataset | 0.380 | 0.006** |
| NIF | Raw KO dataset | Subsampled KO dataset | 0.750 | 0.001*** |

\* $p < 0.05$ ; \*\* $p < 0.01$ ; \*\*\* $p < 0.001$ (significance level of r values)

